# Spatial Transcriptomics Identifies Immune-Stromal Niches Associated with Cancer in Adult Dermatomyositis

**DOI:** 10.1101/2025.03.19.644147

**Authors:** Ksenia S. Anufrieva, Neda Shahriari, Ce Gao, Rochelle L. Castillo, Jessica Liu, Sean Prell, Shideh Kazerounian, Khashayar Afshari, Anastasia N. Kazakova, Erin Theisen, Teresa Bowman, Avery LaChance, Kimberly Hashemi, Ilya Korsunsky, Mehdi Rashighi, Ruth Ann Vleugels, Kevin Wei

## Abstract

Adult-onset dermatomyositis (DM) is an autoimmune inflammatory myopathy with distinct cutaneous manifestations and a strong malignancy association. Through comparative analysis with cutaneous lupus erythematosus (CLE), our integrated spatial and single-cell transcriptomics analysis revealed unique immune and stromal niches associated with DM subtypes. Unexpectedly, we found an association between cancer-associated DM skin lesions and the presence of dispersed immune infiltrates enriched with macrophages, CD8+ T cells, plasma cells, and B cells with preserved vascular architecture. In contrast, non-cancer associated DM skin exhibited dense myeloid cell infiltrates, including neutrophils, monocytes, and macrophages, with elevated expression of IL1B and CXCL10 localized near injured vascular endothelia. Cytokines produced by these myeloid infiltrates together with local tissue hypoxia triggered dramatic stromal remodeling, leading to loss of vascular-associated fibroblasts. In addition to the CXCL10+ myeloid signature, non-cancer-associated DM skin with pDC presence showed the emergence of specific cellular pairs: PD-L1-expressing mregDCs and activated Tregs expressing NFKB2 and TNF receptors. While both DM and CLE showed strong interferon signatures, DM uniquely displayed IFN-β expression. Together, our study provides the first comprehensive spatial mapping of immune and stromal cells in adult-onset DM.

## INTRODUCTION

Adult-onset dermatomyositis (DM) is a rare autoimmune disease characterized by chronic inflammation affecting the skin, muscle, and lungs.^1^ Characteristic cutaneous manifestations of the disease include heliotrope rash, Gottron’s papules and sign, holster sign, shawl sign, V-neck sign and periungual changes including dilated capillary loops and ragged cuticles.^2^ A critical association with DM with onset in adulthood is malignancy^3^, with previous studies demonstrating a lifetime risk of 7% to 30%^4^ and cancer diagnosis occurring either before, concurrent with, or after DM onset.^5^ Malignancy most commonly manifests within the first year of diagnosis but the risk remains elevated beyond that.^5^ Remarkably, adult-onset DM is associated with a wide range of malignancy types, most commonly ovarian, breast, lung, and hematologic cancers, necessitating and extensive and often costly screening workup.^6,7^ While autoantibodies against transcriptional intermediary factor-γ (TIF1γ) and nuclear matrix protein-2 (NXP-2), are associated with increased cancer risk in adult DM patients^8^, the molecular and cellular features underlying cancer association in DM remains incompletely understood.

DM skin pathology is characterized by vacuolar interface dermatitis, increased lymphocytic infiltrate near vasculature, enhanced mucin deposition in the dermis, and basement membrane thickening.^9^ While these histologic findings establish the fundamental pathologic features of DM, they provide limited insight into the complex cellular dynamics of disease pathology and progression, such as the precise organization of immune infiltrates, their molecular programs, and their spatial relationship with surrounding vascular structures. Recent advances in single-cell and spatial transcriptomics have revolutionized our ability to describe complex tissue environments.^10^ These technologies enable comprehensive profiling of cellular ecosystems while preserving crucial information about spatial organization and cell-cell interactions, providing groundbreaking insights into the molecular and cellular mechanisms underlying DM pathogenesis.

In this study, we conducted the first comprehensive integrated analysis of single-cell and spatial transcriptomics of adult-onset DM skin, profiling 44 patient skin biopsies with single-cell RNA sequencing and 17 samples with high-resolution spatial transcriptomics. By comparing DM lesions with both healthy skin and cutaneous lupus erythematosus (CLE), another interferon-driven autoimmune disease with identical histopathological features, we characterized disease-specific cellular states and molecular pathways. Our analysis not only identified features distinguishing DM and CLE from healthy tissue but also stratified DM patients into cancer-associated and non-cancer subgroups. Through spatial transcriptomics, we mapped the architecture of immune cell infiltrates and their interactions with stromal populations in lesional skin, providing a new framework for understanding DM pathogenesis with implications for disease stratification and treatment based on skin biopsy analysis.

## RESULTS

### High-resolution spatial and single-cell atlas reveals interferon-driven tissue reorganization in lesional skin biopsies from DM and CLE

To comprehensively characterize skin pathology in DM and CLE, we conducted high-resolution spatial and single-cell analysis. Using the Chromium Single Cell Gene Expression Flex, we profiled single-cell transcriptomes of 44 samples, encompassing 22 DM patients, 9 CLE patients, 8 non-lesional samples from a subset of these patients, and 6 healthy controls (**Fig. 1A, Extended Data Table 1**). To further delineate the spatial architecture and cell-cell interactions within the tissue microenvironment, we performed high-resolution spatial transcriptomics (Xenium Prime 5K, 10X Genomics) on a subset of biopsies (n=12 lesional DM, n=2 lesional CLE, and n=3 healthy controls), with several patients contributing samples to both the single-cell and spatial transcriptomics analyses to enable direct comparative studies(**Fig. 1A, Extended Data Table 1**). Unlike previous studies^11^ that either employed flow cytometry enrichment or analyzed the dermal and epidermal components of the skin separately, our methodology utilizing intact FFPE tissue sections enabled comprehensive profiling of the complete cellular ecosystem in its native context, including traditionally challenging-to-capture cell states in tissue such as neutrophils and mast cells. Through integrated analysis of both technologies, we identified 16 major skin cell types that were consistently detected across both platforms (**Fig. 1B,C**), with detailed annotation methodology described in the Methods section.

**Fig. 1:**
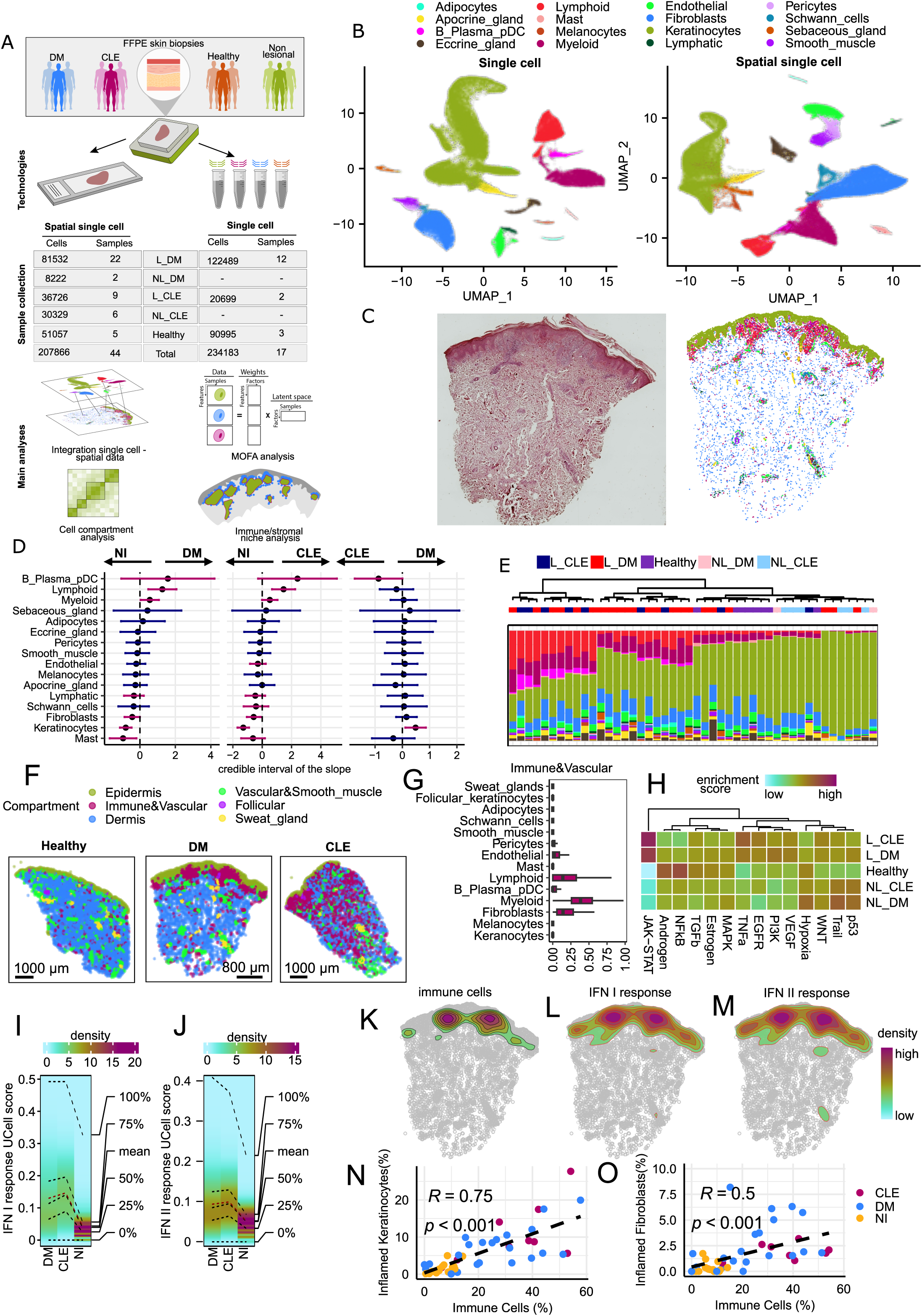
Spatial and single-cell analysis reveals similar patterns of skin remodeling under the action of interferon in OM and CLE. **A.** Overview of experimental design and analysis pipeline. FFPE skin biopsies were analyzed using two complementary approaches: probe-based single-cell transcriptomics (n=44 samples) and spatial transcriptomics (n=18 samples). **B.** UMAP visualization of major cell states identified by single-cell (left) and spatial transcriptomics (right). Each point represents a single cell, colored by cell state name. **C.** Spatial localization of major cell states. The left panel shows H&E staining of an FFPE skin section. Right panel displays spatial distribution of cell states identified by analyzing spatial transcriptomics data at the same FFPE section, with colors matching the cell state legend in Fig. 1B. **D.** Differential composition analysis of major cell states between disease conditions. Points represent the mean of the posterior distribution for the composition parameter, with error bars indicating credible intervals. Red indicates significant changes (FDR < 0.1) and blue shows non-significant changes in cell type proportions. Comparisons are shown for: DM lesional vs non-inflamed (NI) tissue (left), CLE lesional vs NI tissue (middle), and DM lesional vs CLE lesional (right). Arrows indicate which condition shows enrichment for each cell state in the respective comparison. **E.** Major cell state composition of individual samples analyzed by probe-based single-cell transcriptomics. Each row represents a sample, with colors matching the cell state legend in Fig. 1B. Group colors indicate condition: lesional CLE (L_CLE), non-lesional CLE (NL_CLE), lesional DM (L_DM), non-lesional DM (NL_DM), and healthy controls (Healthy). The dendrogram shows hierarchical clustering of samples based on their major cell state composition. **F.** Representative samples from healthy, DM, and CLE showing the spatial distribution of six identified compartments by hoodscanR. **G.** Probability of spatial colocalization between immune&vascular compartment and major cell types. X-axis shows the probability score (0-1.00), with error bars representing the confidence intervals. **H.** PROGENy pathway analysis across different skin conditions. Heatmap shows enrichment scores for pathways across lesional CLE (L_CLE), lesional DM (L_DM), healthy controls (H), and non-lesional samples (NL_CLE, NL_DM). Color intensity represents an enrichment score from low (cyan) to high (red). **I-J.** Density of interferon response scores calculated using UCell for all cells from all samples obtained through probe-based single-cell transcriptomics. Type I (I) and type II (J) interferon response score distributions are shown for DM, CLE, and NI. Color gradient represents cell density, with percentile lines indicating score distribution. **K-M.** Spatial distribution of immune cells and interferon response gene signature in representative lesional DM sample. (K) Kernel density map of immune cells with contour lines highlighting regions of high immune cell concentration. Kernel density maps showing the spatial distribution of cells with above-mean scores for type I (L) and type II (M) interferon response. Color scale indicates density from low (cyan) to high (red). **N-O.** Correlation analysis between immune cell proportion in each sample and CXCL10-expressing populations obtained through probe-based single-cell transcriptomics. (N) Correlation between the percentage of CXCL10+ keratinocytes and immune cell proportion. (O) Correlation between the percentage of CXCL10+ fibroblasts and immune cell proportion. Samples are colored by condition: DM (blue), CLE(red), and NI(yellow). Dashed lines represent linear regression, with R and p-values indicated.

To quantify differences in cellular composition between disease states, we employed sccomp^12^, a computational method designed for differential abundance analysis in single-cell data. Comparison of lesional samples (both DM and CLE) with non-inflamed (NI) tissues (healthy and non-lesional) revealed a shared pattern of immune infiltration in both diseases(**Fig. 1D,E**). Both DM and CLE tissues showed significant enrichment of diverse immune populations, including B cells, plasma cells, and various myeloid and lymphoid cell states (**Fig. 1D,E**). To map the spatial organization of major cell states within the tissue architecture, we employed hoodscanR^13^, a computational tool that uses approximate nearest neighbor algorithm and probability-based clustering to identify cellular neighborhoods in spatial transcriptomics data (**Fig. 1F**). After separating keratinocytes into follicular and interfollicular populations to better distinguish skin regions (**Fig. 1F**, **Extended Data Fig. 1A**), this analysis revealed six distinct tissue compartments: 1) epidermal compartment, primarily composed of interfollicular keratinocytes; 2) immune-vascular compartment, containing B cells, plasma cells, pDCs, lymphocytes, myeloid cells, and endothelial cells; 3) dermal compartment, consisting of fibroblasts and myeloid cells; 4) follicular compartment, containing follicular keratinocytes; 5) sweat gland compartment, containing sweat gland cells; and 6) vascular-smooth muscle compartment, comprising endothelial cells, smooth muscle cells, pericytes, fibroblasts, and myeloid cells. The immune-vascular compartment occupied a substantial area only in lesional samples, predominantly in superficial dermis and localizing near the epidermal compartment (**Fig. 1 F-G**). This pattern of immune cell infiltration is observed in both CLE and DM, consistent with well-documented histopathological findings in DM and CLE^14^. Pathway analysis using PROGENy gene signatures across all conditions revealed significant activation of the JAK-STAT pathway in both DM and CLE lesional skin compared to healthy and non-lesional skin (**Fig. 1H**). Since JAK-STAT signaling is downstream of interferon receptor signaling, we analyzed type I and II interferon response gene sets from the Reactome database using UCell (**Fig.1 I-J**). While both diseases exhibited strong interferon signatures, CLE samples showed higher interferon response scores compared to DM samples. However, we hypothesized that this difference might be attributed to varying levels of immune cell infiltration in our cohort. Indeed, our spatial analysis demonstrated that elevated interferon signaling was predominantly localized to regions of immune cell accumulation, indicating a direct relationship between local immune cell infiltration and inflammatory response (**Fig. 1 K-M**).

Fine-grained analysis of major cell states, including keratinocytes **(Extended Data Fig. 1B-E**) and fibroblasts (**Extended Data Fig. 1H-K**), revealed significant enrichment of interferon-response states (CXCL10_basal, CXCL10_spinous, CXCL10_fib) in lesional samples from DM and CLE compared to non-inflamed tissues (**Extended Data Fig. 1C, 1I**). These interferon-response clusters were characterized by high expression of chemokines CXCL10, CXCL11, and CXCL9 (**Extended Data Fig. 1D, 1J).** Moreover, the relative abundance of CXCL10+ cell states within both keratinocytes and fibroblasts showed strong positive correlation with the proportion of immune cells in the tissue (**Fig. 1N-O**). Among keratinocytes, we identified an expanded population of proliferative basal keratinocytes (cyclic_basal) in lesional samples that express both proliferation markers (MKI67) and DNA damage-response genes (FANCA, FANCI, FOXM1, CHEK1)(**Extended Data Fig. 1C-E**). Additionally, both DM and CLE showed increased proportions of granular keratinocytes expressing high levels of CALM3 and CALM5 (members of the calmodulin gene family that encode calcium-binding proteins essential for calcium homeostasis)(**Extended Data Fig. 1C-E**). Detailed characterization of fibroblast heterogeneity revealed that beyond CXCL10-expressing fibroblasts, lesional skin in both conditions showed elevated numbers of CCL19-expressing fibroblasts (CCL19_fib) and interferon-associated IFIT2-expressing fibroblasts (IFIT2_fib)(**Extended Data Fig. 1H-J**).

### Single-cell analysis reveals immune cell heterogeneity in DM and CLE

Since immune infiltration in lesional skin is a hallmark feature of both DM and CLE, we queried our single-cell data to identify immune cell composition with disease-specific association. We first identified fine-grained cell states for lymphoid cells, myeloid cells, B cells, and plasma cells, with the detailed annotation process described in the Methods section(**Fig. 2A, Extended Data Fig. 2A-C**). Analysis of immune cell composition among all lesional skin using Bray-Curtis dissimilarities revealed distinct patterns of immune cell infiltration (**Fig. 2B-C**). The proportion of immune cells relative to total cells emerged as the primary factor differentiating skin biopsies, accounting for approximately 45% of the variance in cellular composition (PCoA1, **Fig. 2B**). Surprisingly, the second most significant distinguishing factor was the presence or absence of cancer in DM patients, explaining approximately 15% of the variance (PCoA2, **Fig. 2C**). To identify disease-specific immune signatures, we performed differential abundance analysis using sccomp to compare immune cell populations across CLE, cancer-associated DM, and non-cancer DM skin (**Fig. 2D, Extended Data Fig. 2D**). This analysis revealed that non-cancer associated DM skin exhibited a distinctly different immune composition compared to both CLE and cancer DM skin. Non-cancer associated DM showed increased proportions of mature regulatory dendritic cells (mregDCs, BIRC3+LAMP3+), neutrophils, monocytes, CXCL10+ myeloid cells, and conventional type 2 dendritic cells (cDC2s). Conversely, these biopsies showed reduced frequencies of B cells, plasma cells, and GZMK+GZMB+ CD8+ T cells compared to cancer DM and CLE biopsies (**Fig. 2D, Extended Data Fig. 2A**). Further analysis revealed several additional disease-specific features. Most notably, we observed a near-complete absence of neutrophils in CLE biopsies identifying neutrophil infiltration as a distinctive feature of DM pathology. CLE samples also showed near-complete depletion of Langerhans cells. Additionally, while CLE samples exhibited significantly lower levels of proliferating lymphocytes, compared to both cancer and non-cancer DM biopsies, they showed increased numbers of plasmacytoid dendritic cells (pDCs) and plasma cells (**Fig. 2D, Extended Data Fig. 2D**).

**Fig. 2:**
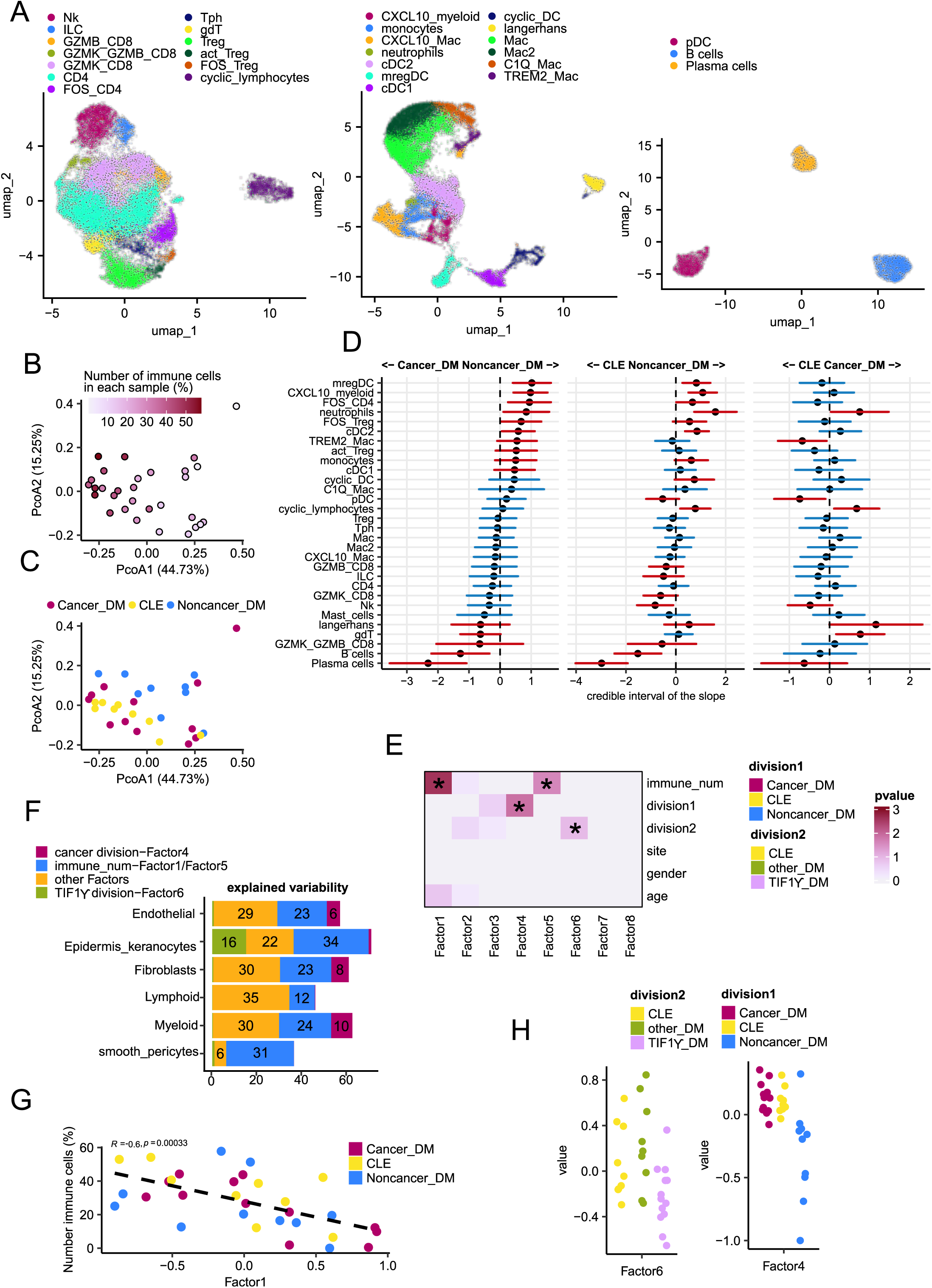
Heterogeneity of immune cells infiltration determines differences in pathologic features of OM and CLE. **A.** UMAP visualization of fine-grained cell state analysis of lymphocytes, myeloid cells, B cells/plasma cells/pDC, colored by cell states identified during the analysis of single-cell data. **B-C.** Principal Coordinates Analysis of Bray-Curtis dissimilarities of immune cell state composition across lesional samples. (B) Colors ranging from pink (low) to red (high) represent the proportion of immune cells relative to total cells in each lesional sample. (C) Colors represent clinical information: cancer-associated DM, CLE, and non-cancer-associated DM lesional samples. **D.** Differential composition analysis of immune cell states across disease conditions in lesional tissue. Points show the mean posterior distribution of composition differences, with error bars representing credible intervals (2.5% and 97.5% quantiles). Red indicates significant changes (FDR < 0.1) and blue shows non-significant differences in cell state proportions. Comparisons are shown between: Cancer_DM vs. Noncancer_DM (left), CLE vs. Noncancer_DM (middle), and CLE vs. Cancer_DM (right). Arrows indicate which condition shows enrichment for each cell state in the respective comparison. **E.** The heatmap shows -log10 transformed adjusted p-values from Kruskal-Wallis tests examining relationships between MOFA factor scores (x-axis) and clinical variables (y-axis). Darker colors indicate stronger associations, and asterisks (*) denote statistically significant associations. **F.** The stacked bars represent the percentage of gene expression variance explained by different MOFA factors for each cell type. Cell types are shown on the y-axis, with the percentage of explained variance on the x-axis. **G.** The x-axis shows MOFA Factor 1 scores, and the y-axis shows the percentage of immune cells relative to total cell number in each sample. Points are colored by disease type: Cancer_DM (red), CLE (yellow), and Noncancer_DM (blue). The dashed black line shows correlation between Factor 1 and immune cell proportion). **H.** Distribution of MOFA factor scores across disease subtypes. The left panel displays Factor 6 stratification by disease and TIF1γ status: TIF1γ-positive DM (purple), TIF1γ-negative DM (green), and CLE (yellow). The right panel shows Factor 4 stratification by disease and cancer status: Cancer-associated DM (red), Non-cancer DM (blue), and CLE (yellow). Each dot represents a patient sample.

### Multicellular factor analysis uncovers distinct stromal and immune cell signatures in non-cancer associated DM skin

The striking immunological differences between cancer-associated and non-cancer DM skin (**Fig. 2B-D**) led us to investigate transcriptional programs underlying cancer-association across all skin cell types. We employed Multicellular Factor Analysis (MOFA)^15^, which provides an unsupervised approach to studying transcriptional programs across all cell populations in disease states at the same time.

Our MOFA model effectively captured the majority of gene expression variability across all studied cell types constructing a new latent space composed of eight factors (**Fig. 2E**). Factor 1 emerged as the primary driver of gene variation, explaining variability across all cell populations and containing numerous interferon response genes. Kruskal-Wallis analysis of the MOFA factors revealed its strong association with immune infiltration percentage (fdr < 0.00261)(**Fig. 2E-G**). Unexpectedly, we found factor 4 to be strongly associated with the distinction between cancer-associated DM, non-cancer DM, and CLE samples (fdr < 0.0141)(**Fig. 2E**). Factor 4 explained cell type-specific gene expression variation, primarily in myeloid cells and fibroblasts (8% each) and endothelial cells (6%)(**Fig. 2E,H**). Interestingly, the expression of ABCA8, ABCA9, and ABCA10 genes within the fibroblasts emerged as the primary drivers of Factor 4 (**Extended Data Fig. 2E**). Factor 6 uniquely captured keratinocyte-specific variation (16%) and was significantly associated with TIF1γ-positive DM status (pvalue < 0.0152, fdr < 0.121)(**Fig. 2E,H**).

### Unique spatial organization of immune cell niches reveals disease-specific archetypes in DM and CLE lesions

Our spatial transcriptomics data demonstrated that infiltrating immune cells form structured aggregates rather than random distributions, particularly concentrating in superficial dermis near interfollicular keratinocytes and vascular structures. (**Extended Data Fig. 3A**). To characterize these immune niches and identify disease-specific patterns, we implemented a computational pipeline consisting of two key steps: (1) HDBSCAN clustering to identify spatially distinct immune cell niches based on their physical proximity, and (2) Bray-Curtis dissimilarity analysis to evaluate the compositional differences between these niches across disease conditions (**Fig. 3A, Extended Data Fig. 3A**). This approach enabled us to systematically map and characterize the cellular compositions of immune niche organization across different disease states (**Fig. 3B**).

**Fig. 3:**
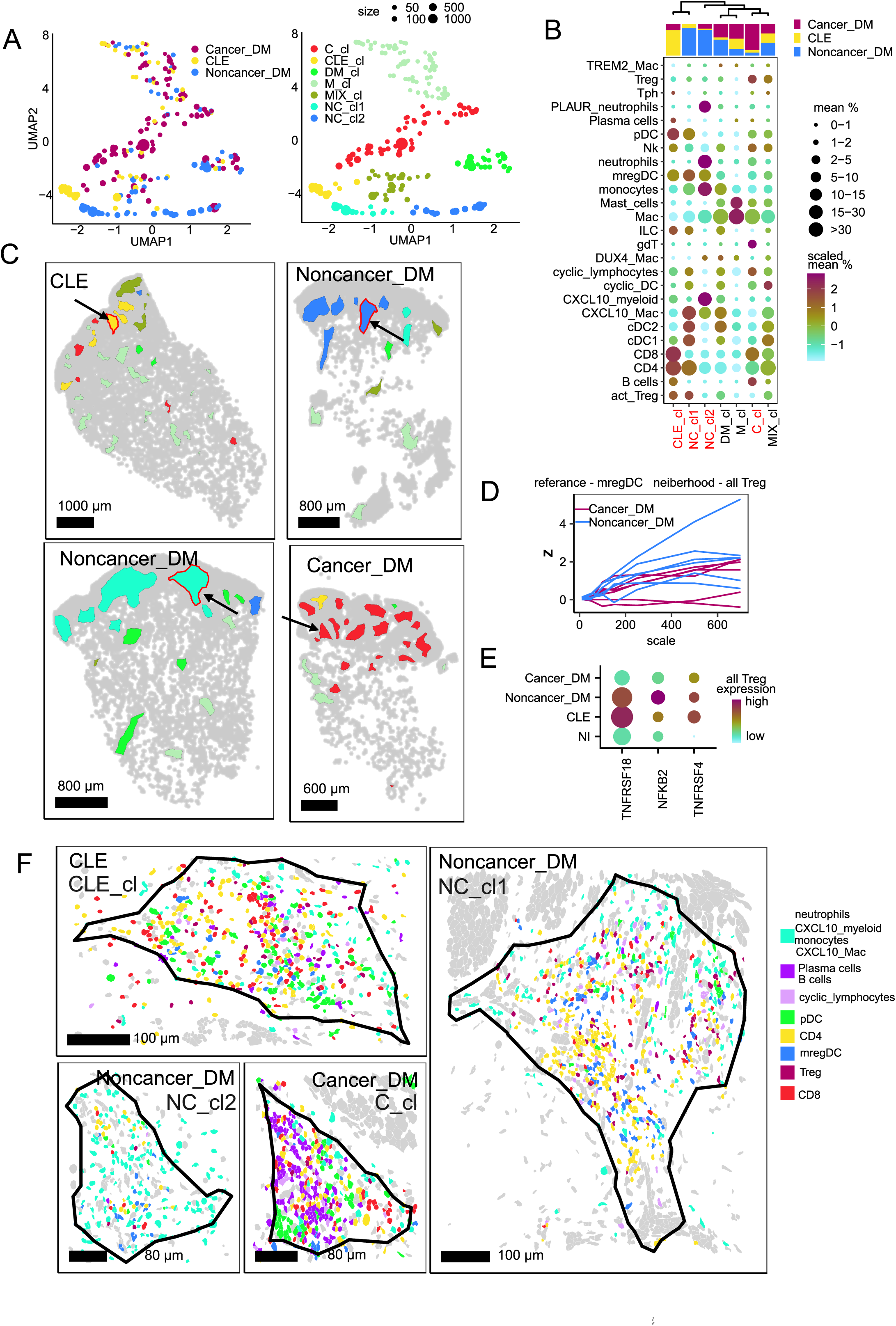
Spatial organized immune niches define cancer and non-cancer associated OM. **A.** UMAP visualization of immune cell composition within spatially defined immune niches from 14 lesional samples obtained through analysis of spatial single cell data. (A) Left panel shows immune niches colored by disease condition: Cancer_DM (red), CLE (yellow), and Noncancer_DM (blue). Right panel displays the same aggregates colored by niche type (C_cl, CLE_cl, DM_cl, M_cl, MIX_cl, NC_cl1, NC_cl2). Point size represents the number of immune cells in each niche. **B.** Immune cell composition of identified immune niche types. Dot plot showing the distribution of immune cell states (y-axis) across different immune niches (x-axis). Dot size represents the mean percentage of each cell state within the niche. Color scale shows the relative abundance of each cell state across conditions, from high (red) to low (cyan). Top bar indicates the disease condition associated with each immune niche type: Cancer_DM (red), CLE (yellow), and Noncancer_DM (blue). **C.** Spatial location of immune niches in representative skin sections from CLE, Non-cancer DM, and Cancer-associated DM lesional samples. Colored regions represent different immune niche types as defined in Fig. 3A (C_cl, CLE_cl, DM_cl, M_cl, MIX_cl, NC_cl1, NC_cl2), while cells outside immune niches are represented in gray. **D.** Spatial colocalization analysis between mregDCs (reference) and Tregs (neighbors). The plot shows enrichment Z-scores (y-axis) calculated at increasing spatial distances in micrometers (x-axis). Lines represent individual samples from Cancer_DM (red) and Noncancer_DM (blue), with higher Z-scores indicating stronger spatial colocalization than expected by chance. **E.** Expression of activated Treg gene signature across different disease conditions. Dot plot shows single-cell gene expression data for TNFRSF18, NFKB2, and TNFRSF4 in all Treg cells from Cancer_DM, Noncancer_DM, CLE, and NI samples. **F.** Spatial organization within representative immune niches from different disease conditions. Each chosen niche is highlighted with an arrow in Fig. 3B. Each immune cell type is represented by a distinct color as shown in the legend. Black outlines display the boundaries of immune niches.

Analysis of immune niches across 14 lesional samples revealed two major niche archetypes: low-density niches (M_cl, C_cl, DM_cl, MIX_cl) and high-density niches (CLE_cl, NC_cl1, NC_cl2) (**Fig. 3A-C, Extended Data Fig. 3B-C**). Niche subtypes were named according to their disease association: NC for non-cancer DM-associated niches, CLE for cutaneous lupus-specific niches, C for cancer-associated DM niches, DM for niches shared between cancer and non-cancer DM, MIX for non-disease-specific niches, and M for macrophage-enriched niches. Low-density niches displayed relatively sparse distribution with reduced cell density and were highly occupied by macrophages (**Extended Data Fig. 3B-C**). Within the low-density niche archetype, M_cl niche subtype showed enrichment for both macrophages and mast cells, localizing predominantly to reticular and deep dermal areas (**Extended Data Fig. 3D-F**). This niche subtype is observed in both CLE and DM, suggesting a shared inflammatory response across CLE and DM. The second low-density niche subtype, C_cl, displayed a unique disease-specific pattern, being predominantly present in cancer-associated DM (**Fig. 3A-C, Extended Data Fig. 3D-F**). This subtype contained substantial populations of macrophages, CD8+ (∼12%) and CD4+ T cells (∼11%), and notably elevated levels of γδT cells, B cells, plasma cells compared to other niche subtypes (**Fig. 3B,C,F, Extended Data Fig. 3E,G**).

In contrast to low-density niches, high-density niches displayed compacted structures with substantial immune infiltration. Among high-density niche archetypes, we identified two niche subtypes (NC_cl1, NC_cl2) uniquely associated with non-cancer-DM that were characterized by high proportions of CXCL10-expressing myeloid cells such as neutrophils, monocytes, and CXCL10+ macrophages. NC_cl1 showed enrichment in CXCL10+ macrophages, CD4+ T cells, activated Tregs, mregDC, cDC2, and pDCs, while NC_cl2 featured elevated neutrophils, monocytes, and mregDC (**Fig. 3B,C,F**).

Compositional analysis revealed that beyond CXCL10+ myeloid cell enrichment in non-cancer DM, a key distinction from cancer-associated DM was the increase in abundance and co-localization of Tregs and mregDCs in non-cancer DM (**Fig. 3 B,D**). Furthermore, Tregs in non-cancer DM showed elevated expression of TNFRSF18, TNFRSF4, NFKB2, and NFKBIA, with activated Tregs present only in non-cancer-associated immune niches (NC_cl1), suggesting Treg activation via mregDCs within this niche (**Fig. 3 B,E**).

The CLE-specific cluster (CLE_cl) displayed a unique immune composition combining features of both non-cancer DM and cancer-associated DM clusters (**Fig. 3B,F, Extended Data Fig. 3F**). Compared to DM-associated immune niches, CLE_cl was distinctly enriched for pDCs and peripheral T helper cells (Tph cells)(**Fig. 3B, Extended Data Fig. 3F**). Like NC_cl1, it contained high levels of CD4+ T cells, activated Tregs, mregDCs, and CXCL10+ macrophages, while sharing characteristics with C_cl through elevated CD8+ T cells, plasma cells, and B cells. Consistent with our single-cell analysis, CLE immune aggregates lacked neutrophils and showed a reduced number of lymphocytes with elevated proliferation gene signature.

### Disease-associated transcriptional programs define distinct immune niches

To characterize molecular programs within immune cell niches, we performed pseudobulk analysis of immune niches from lesional biopsies. Principal component analysis revealed robust separation between conditions, with PC1 (25% variance) distinguishing DM from CLE immune niches (**Extended Data Fig. 4A**). Differential expression analysis across CLE, non-cancer-associated DM, and cancer-associated DM lesions identified distinctive immune signatures (**Fig. 4A**). For example, non-cancer-associated DM immune niches showed significant upregulation of pro-inflammatory mediators specifically expressed by neutrophils, monocytes, and CXCL10+ myeloid cells, including inflammatory calcium-binding proteins S100A8/S100A9, pro-inflammatory cytokine IL1B, monocyte chemoattractant CCL2, and T-cell recruiting chemokine CXCL10 (**Fig. 4A-B**).

**Fig. 4:**
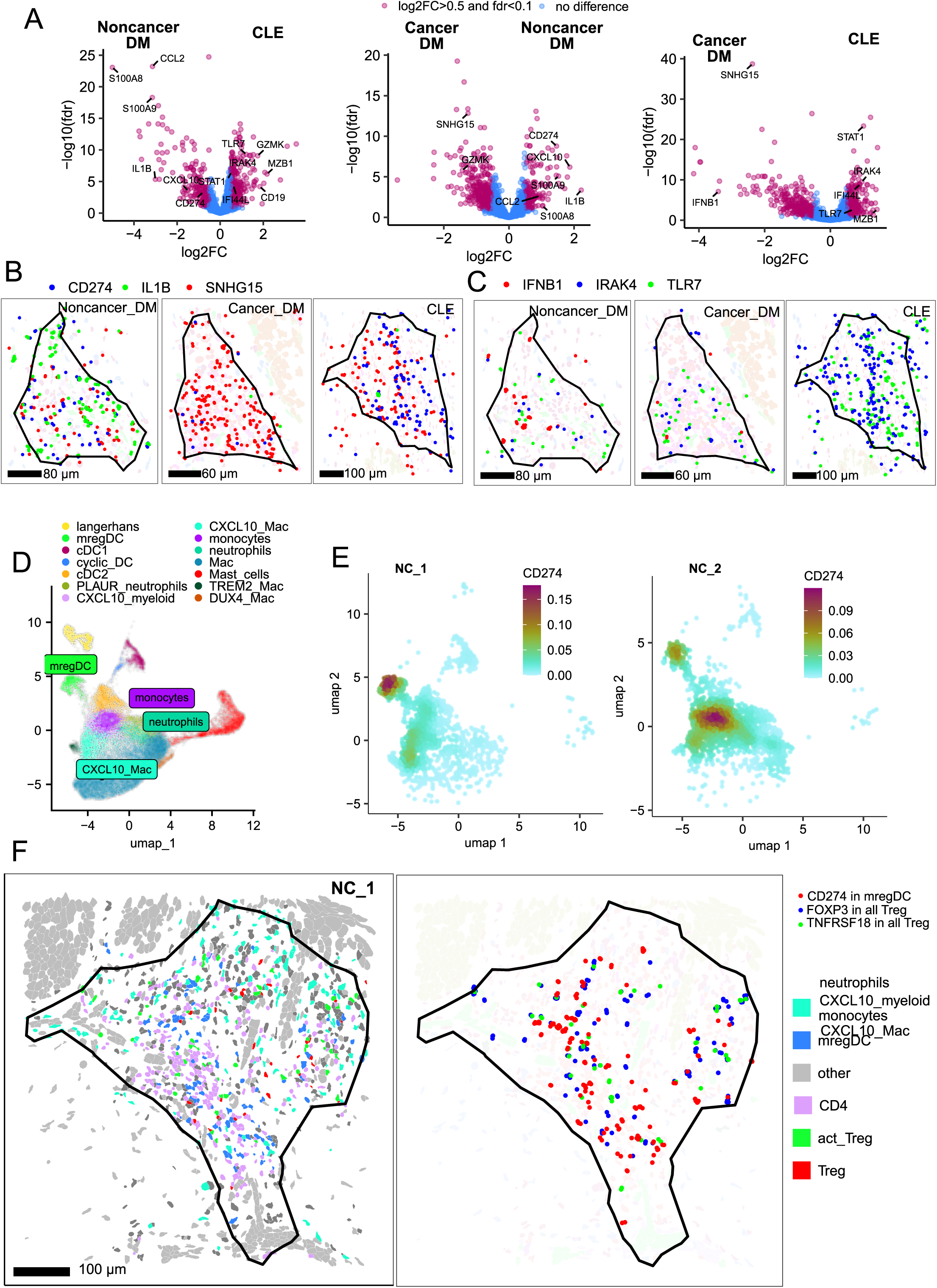
Activated Treg and PD-L1+ mregDC distinguish immune niches in non-cancer assosiated OM. **A.** Differential gene expression analysis of immune niches between different disease conditions. Volcano plots show gene expression differences: Noncancer_DM vs CLE (left), Cancer_DM vs Noncancer_DM (middle), and Cancer_DM vs CLE (right). Red dots indicate significantly differentially expressed genes (log2FC > 0.5 and FDR < 0.1), while blue dots represent non-significant changes. Key disease-associated genes are labeled. **B-C.** Spatial distribution of key differentially expressed genes in representative immune niches from each disease condition. (B) Distribution of CD274 (blue), IL1B (green), and SNHG15 (red) transcripts within immune niches. (C) Distribution of IFNB1 (red), IRAK4 (blue), and TLR7 (green) transcripts within immune niches. **D.** UMAP visualization of myeloid cell states identified through analysis of spatial transcriptomics. Each point represents a single cell, colored by cell type. **E.** UMAP visualization showing density of CD274-expressing cells from the NC_1 and NC_2 niche. Cell positions correspond to the myeloid cell state UMAP embeddings shown in Fig. 4D. **F.** Spatial localization of mregDC and Treg cell interactions in NC_cl1 immune niches. The left panel shows the distribution of immune cell state, colored according to the legend. Right panel displays the spatial distribution of specific transcripts: CD274 in mregDC (red), FOXP3 in all Tregs (blue), and TNFRSF18 in all Tregs (green), indicating potential functional interactions between these cell states. The niche shown is the same as in Fig. 3F, with black outlines delineating niche boundaries.

We found elevated CD274 expression, encoding PD-L1, in non-cancer-associated DM and CLE immune niches compared to cancer-associated DM samples (**Fig. 4A-B**). While CD274 expression was restricted to mast cells in healthy skin, lesional skin showed broad expression across mregDCs, CXCL10+ myeloid cells, neutrophils, CXCL10+ macrophages, and monocytes, reflecting its critical role in immune regulation.^16^ Notably, CD274 expression patterns varied across non-cancer-associated DM niches. NC_cl2 aggregates showed expression limited to CXCL10+ myeloid populations (**Fig. 4D-E**), while NC_cl1 niches displayed CD274 expression in both mregDCs and CXCL10+ myeloid cells (**Fig. 4D-E**). Importantly, CD274 expression in mregDCs coincided with activated Treg signatures (**Fig. 4F**), whereas CD247 expression in CXCL10+ myeloid cells did not spatially coincide with an activation phenotype in Tregs (**Extended Data Fig. 4B**).

Also, our spatial transcriptomics data revealed differential expression of IFNβ within immune niches, with higher levels observed in both cancer and noncancer-associated DM compared to CLE immune niches (**Extended Data Fig. 4C**, **Fig. 4C**), supporting a critical role of IFN-B in DM pathology.^17,18^ Furthermore, single-cell analysis of DM and CLE samples confirmed IFNβ expression specifically in myeloid cells in 3 DM samples, suggesting that presence of IFNβ expression may distingusih DM from CLE. Due to the low expression level of certain cytokines, both single-cell and spatial single-cell sequencing have limited ability to assess the abundance of critical immune mediators such as IL2, IL10, and IFNα.

Disease-specific signatures were further characterized by elevated expression of interferon response genes (STAT1, IFI44) and innate immune components (IRAK4, TLR7) in CLE samples (**Extended Data Fig. 4A**, **Fig. 4A,C**), consistent with lupus pathogenesis.^19,20^ Cancer-associated DM samples uniquely showed upregulation of cancer-associated genes, including SNHG15 (**Extended Data Fig. 4D**, **Fig. 4B**), a lncRNA frequently elevated across multiple malignancies.^21,22^

### Vascular remodeling and metabolic reprogramming underlie distinct immune niche phenotypes

Building on our observation that immune cells organize into distinct niches centered around skin vasculature, we next characterized endothelial and fibroblasts populations within immune niches and their surrounding areas, as defined within 50 micrometers of niche boundaries (**Fig. 5A**). Using Bray-Curtis dissimilarity analysis of endothelial cells within and adjacent to immune niches we identified four distinct vascular niches (**Fig. 5B-C, Extended Data Fig. 5 A-C**). Two niches showed no disease-specific associations: the lymphatic_cl (47% lymphatic cells) and the DLL4_cl (49% DLL4+ cells) (**Fig. 5C, Extended Data Fig. 5D-F**). The DLL4_cl niche was predominantly located in reticular dermis and was strongly associated with non-inflammatory macrophage-enriched immune niches (**Extended Data Fig. 5C-F**). In contrast, we identified two endothelial niches with disease-specific associations. The CXCL10_cl niche, predominantly found in CLE and non-cancer associated DM samples, comprised CXCL10-positive venous (34%) and CXCL10-positive capillary cells (8%) (**Fig. 5C, Extended Data Fig. 5D-E**). Similar analysis of fibroblast populations within and adjacent to immune niches also revealed increased abundance of CXCL10-expressing cell states in non-cancer DM and CLE samples compared to cancer-associated DM (**Extended Data Fig. 5G-H)**. The venous_cl niche, enriched in cancer-associated DM samples, consisted primarily of venous endothelial cells (49%) lacking both CXCL10 and DLL4 signatures (**Fig. 5C, Supplementary** Fig. 5F). Notably, non-cancer DM samples displayed a more inflammatory signature in both capillary and venous populations compared to cancer DM. Furthermore, endothelial vasculature in cancer DM samples appeared more rounded with open lumens(**Fig. 5D**). To quantify vascular alterations in DM skin lesions, we measured endothelial cell density within 600-μm regions extending from the upper edge of keratinocytes through the papillary dermis (**Fig. 5F)**. Quantitative analysis revealed significantly higher endothelial cell density in both cancer and non-cancer-associated DM compared to CLE, corroborating previous studies documenting increased vascularization in DM lesions **(Fig. 5G**)^23^. Within these regions, cancer-associated DM samples in particular showed elevated endothelial cell density with a marked increase in endothelial tip cells - endothelial cells that guide vessel formation during sprouting angiogenesis (**Fig. 5H**). Consistent with productive sprouting angiogenesis in cancer-associated DM, our compositional analysis showed enrichment of venous cells in cancer-associated DM lacking both CXCL10 and DLL4 signature, while non-cancer DM and CLE samples displayed elevated CXCL10-positive capillary and venous cells (**Fig. 5H**).

**Fig. 5:**
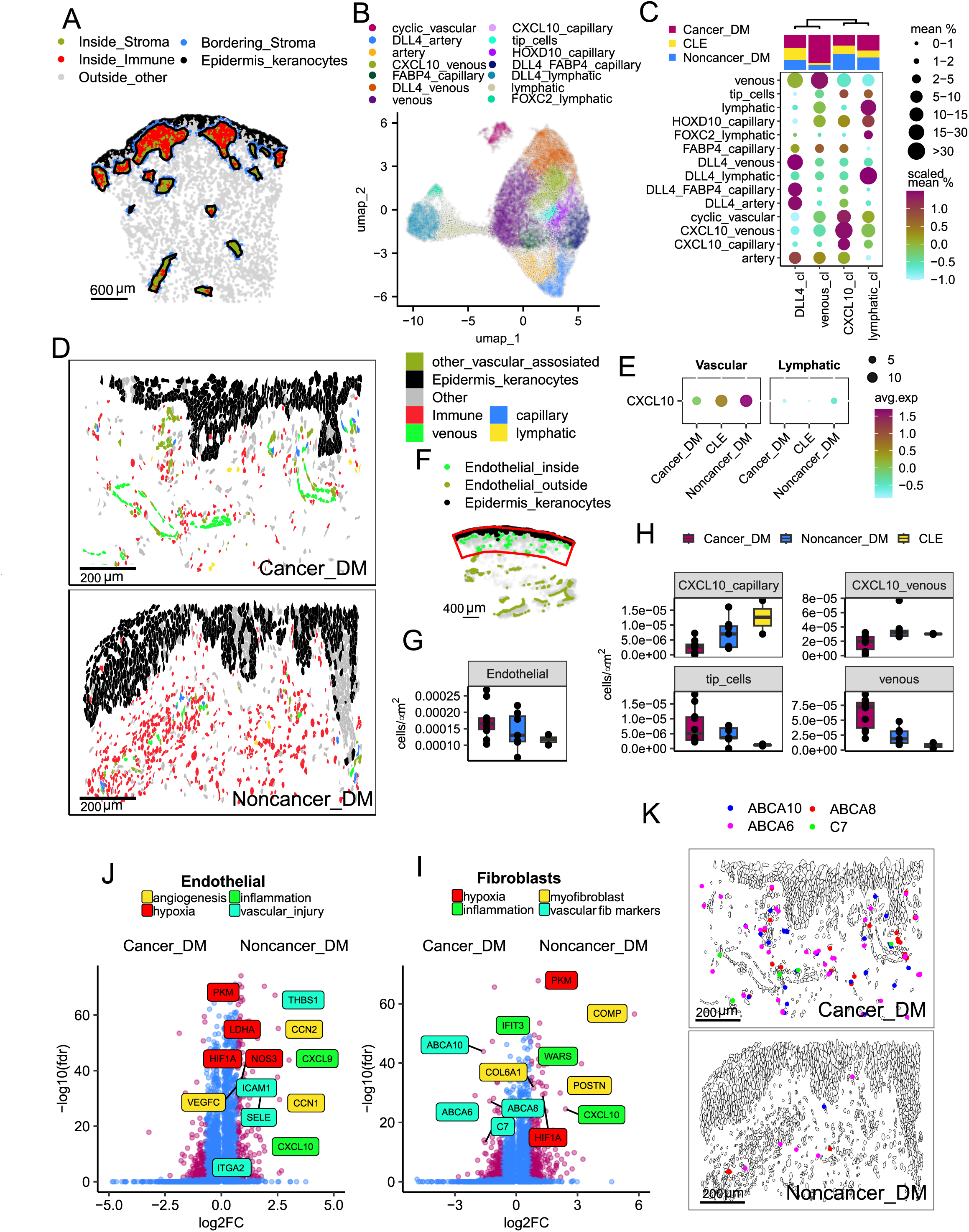
Disease-specific vascular and stromal niches are shaped by hypoxia and inflammation. **A.** Pipeline for stromal analysis within immune niches. Representative skin section showing distinct spatial regions: immune cells (red), stromal cells within immune aggregates (green), bordering stromal cells within 50 micrometers of immune aggregates (blue), epidermal keratinocytes (black), and other cells (gray). **B.** UMAP visualization of endothelial cell states identified by analyzing spatial transcriptomics data. Each point represents a single cell, colored by cell state identity as shown in the legend. **C.** Endothelial cell composition within and adjacent to immune niches, grouped into distinct vascular niches. The dot plot displays endothelial states (y-axis) within each identified vascular niche type, where dot size represents the mean cell type percentage. Color scale indicates the relative abundance of each cell type (red: high, cyan: low). The top bar shows the distribution of samples from different conditions within each niche type: Cancer_DM (red), CLE (yellow), and Noncancer_DM (blue). **D.** Representative spatial distribution of cell states in Cancer_DM and Non-cancer DM samples representing typical endothelial cell structure morphologies between these two conditions. **E.** Expression of CXCL10 in vascular and lymphatic endothelial cells across disease conditions. The dot plot shows normalized and scaled CXCL10 expression levels in vascular (left) and lymphatic (right) endothelial states from Cancer_DM, Noncancer_DM, and CLE samples. Dot size indicates cell number, and color represents average scaled expression **F.** Pipeline for endothelial cell density quantification. Representative skin section demonstrating the selection of standardized analysis regions: epidermal keratinocyte layer (black) is used as a reference to define polygonal regions extending 600 μm into the dermis. Endothelial cells are classified as inside (light green) or outside (dark green) in these regions. **G-H.** Endothelial cell density quantification across disease conditions. (G) Box plots show the number of endothelial cells per square micrometer within standardized 600-μm regions from the epidermal border. (H) Box plots show the number of cells per square micrometer within standardized 600-μm regions from the epidermal border for different endothelial states: CXCL10+ capillary, CXCL10+ venous, tip cells, and venous cells. Samples are colored by disease type: Cancer_DM (red), Noncancer_DM (blue), and CLE (yellow). Each dot represents an individual sample. **J-I.** Differential gene expression analysis in endothelial cells(J)/fibroblasts(I) between Cancer_DM and Noncancer_DM immune niches. Volcano plot shows gene expression differences, with significantly differentially expressed genes (|log2FC| > 0.7, FDR < 0.1; red dots) labeled and colored by pathway for endothelial cells (J): angiogenesis (yellow), hypoxia (red), inflammation (green), and vascular injury (turquoise). For fibroblasts (I): fibrosis (yellow), hypoxia (red), inflammation (green), and vascular-associated signature (turquoise). Blue dots represent non-significant changes. **K.** Spatial distribution of vascular-associated fibroblast gene signature in Cancer_DM and Non-cancer DM samples. Representative tissue sections showing expression of ABCA10 (blue), ABCA8 (red), ABCA6 (pink), and C7 (green) transcripts in fibroblasts. Regions correspond to those shown in Fig.5D.

To integrate immune and stromal analyses, we examined differential gene expression across disease-associated immune niche populations. This analysis revealed that stromal cells within non-cancer-associated niches (NC_cl1, NC_cl2), when compared to cancer-associated aggregates (C_cl1), exhibit a pronounced upregulation of hypoxia- and inflammation-related pathways. Specifically, endothelial cells from NC_cl1 and NC_cl2 niches displayed increased expression of hypoxia-inducible genes (HIF1A, PKM, NOS3, LDHA) and inflammatory mediators (CXCL10, CXCL9, CXCL11) (**Fig. 5J**). Additionally, these endothelial cells exhibited upregulation of genes involved in vascular injury (ICAM1, THBS1, ITGA2, SELE) and angiogenesis (VEGFC, CCN1, CCN2), indicating a coordinated transcriptional response to alteration in oxygen tension within these lesional microenvironment (**Fig. 5J**). Consistently with a regional response to hypoxia, fibroblasts from NC_cl1 and NC_cl2 niches, compared to those in cancer-associated aggregates (C_cl1), exhibited significant upregulation of genes involved in fibrotic pathway genes (POSTN, COMP, COL6A1, COL4A1) along with enhanced transcription of hypoxia (HIF1A, PKM) and inflammation-related genes (CXCL10, CXCL9, CXCL11, IFIT3, WARS, HLA-DRA)(**Fig. 5I**). Notably, fibroblasts in NC_cl1 and NC_cl2 niches displayed a significant downregulation of ABCA10, ABCA8, C7, C3 and ABCA6, with expression levels approaching near-zero—contrasting with cancer-associated fibroblasts (C_cl1) and CLE-associated fibroblasts (CLE_cl)(**Fig. 5J, Extended Data Fig. 5I).** These genes, predominantly expressed in vascular-associated fibroblasts, serve as key molecular markers of this stromal population (**Extended Data Fig. 5J**). Furthermore, analysis of vascular-associated fibroblasts from healthy skin confirmed a high expression of ABCA10, ABCA8, C7, C3,ABCA6, suggesting that their loss in non-cancer immune niches represents a fundamental shift in the stromal microenvironment (**Extended Data Fig. 5K)**.

Since all stromal populations associated with non-cancer-associated immune aggregates exhibit elevated expression of hypoxia- and inflammation-related genes, we performed a spatial correlation analysis to investigate their relationship with other gene expression patterns. Our analysis revealed that angiogenesis, hypoxia, inflammation, and vascular injury signatures were directly correlated with the expression of cytokines highly expressed by monocytes and neutrophils, including CCL2, CCL8, S100A8, S1009, CXCL10, and IL1B (**Fig. 6A,B**). In contrast, the vascular-associated gene signature (ABCA8, ABCA9, C7, C3) displayed an inverse correlation with these cytokines, suggesting that hypoxia and inflammatory signals actively suppress vascular-associated fibroblast transcriptional program (**Fig. 6A,B**), resulting in loss of vascular fibroblast identity and injury to the steady-state perivascular compartment. However, this analysis does not clarify the main cause of the suppression of vascular-associated fibroblasts–whether the drivers are the cytokines highly expressed by monocytes and neutrophils or hypoxia. To address this question, we performed an in vitro experiment using human dermal fibroblast cultures exposed to both inflammatory cytokines and hypoxia. Fibroblasts were cultured under either normoxic or hypoxic (0.5% O₂) conditions and treated with IL1-β (1 ng/mL), IFN-β (5 ng/mL), or IFN-γ (5 ng/mL) for 24 hours. Our qPCR analysis revealed that hypoxia significantly reduced the expression of vascular-associated fibroblast markers (C7, ABCA6, ABCA8) compared to normoxic conditions, regardless of cytokine treatment (**Fig. 6C**). While cytokine treatment had a mild reduction in the expression of vascular fibroblast markers. Together, the dramatic reduction observed under hypoxic conditions suggests that oxygen deprivation is the primary driver of vascular-associated fibroblast identity loss in non-cancer DM skin.

**Fig. 6:**
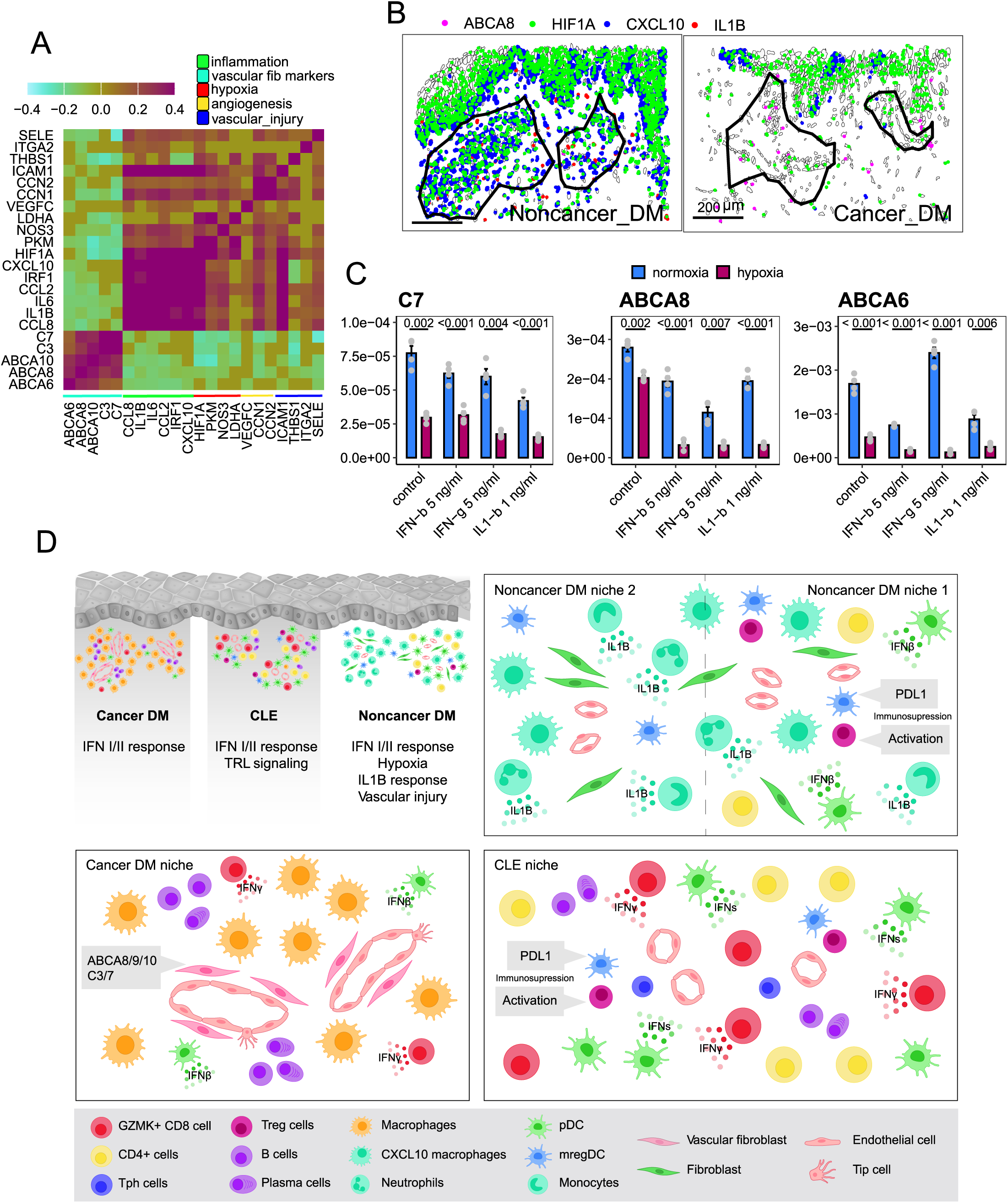
Hypoxia-driven suppression of vascular fibroblast identity and its spatial correlation with inflammatory signals. **A.** Spatial correlation analysis of gene expression within all identified immune niches. Heatmap displaying the correlation coefficients between genes associated with inflammation (green), vascular fibroblast markers (red), hypoxia (purple), angiogenesis (yellow), and vascular injury (blue). Positive correlations are shown in red, while negative correlations appear in cyan. **B.** Spatial distribution of key gene signatures in immune niches across disease conditions. Representative spatial transcriptomic maps showing the distribution of ABCA8 (pink), HIF1A (green), CXCL10 (blue), and IL1B (red) transcripts within Non-cancer DM (left) and Cancer-associated DM (right) immune niches. Black outlines delineate the niche boundaries. Each dot represents a single detected transcript using the spatial transcriptomics platform. **C.** qPCR analysis of vascular fibroblast markers (C7, ABCA6, ABCA8) in dermal fibroblasts treated with IL-1β (1 ng/mL), IFN-γ (5 ng/mL), and IFN-β (5 ng/mL) under normoxic (blue) and hypoxic (purple) conditions for 24 hours. Relative expression levels (normalized to GAPDH)) are shown, with error bars representing standard deviation. Statistical significance was determined using a two-tailed Student’s t-test. *ns* indicates no significant difference. **D.** Schematic representation of immune-stromal organization and molecular signatures in cancer-associated DM, non-cancer DM, and CLE niches.

## DISCUSSION

The pathology of autoimmune skin diseases has traditionally relied on histopathologic assessment of skin biopsies, which identifies features of inflammation, but is insufficient to identify cellular and molecular features specific to distinct inflammatory etiologies.^24,25^ Our comparative analysis of DM and CLE provides several key advantages–both conditions share similar histological features of interface dermatitis and prominent interferon signatures, allowing us to identify truly disease-specific rather than general inflammatory pathways (**Fig. 6D**). Firstly, in our study, we identified DM-specific expression of IFNβ in myeloid cells, while CLE showed broader interferon response through TLR/IRAK signaling. It has been previously shown that IFNβ plays a central role in DM pathogenesis, with elevated plasma levels^26^ strongly correlating with muscle disease activity.^7^ Similarly, studies have observed a specific correlation between cutaneous activity scores and IFNβ, but not IFNα protein levels.^17^ The clinical efficacy of IFNβ inhibition, with dazukibart–an IFNβ specific monoclonal antibody, has shown promise in phase 2 clinical trials for DM treatment.^18^ Our spatial and single cell analysis, in addition, revealed increased pDC abundance in CLE lesions, organized in distinct immune niches with elevated TLR/IRAK signaling. The co-localization of pDCs, known producers of IFNα through TLR7/9 activation,^28^ suggests a coordinated inflammatory program distinct from the IFNβ-driven pathology in DM. Secondly, DM skin biopsies exhibit prominent perturbation in the vascular compartment. Emergence of endothelial tip cells and increased vascular density in DM lesions suggest formation of new blood vessels, or angiogenesis, in some DM lesions. On the other hand, non-cancer-associated DM lesions showed evidence of rarefaction, loss of perivascular fibroblast identity, and regional hypoxia, suggesting early vasculopathy, which is a known feature of DM lesions.^29^ Third, we identified lesional neutrophil infiltration as a DM-specific feature and absent in CLE lesions. Studies have shown neutrophils and their extracellular traps (NETs) correlate with interstitial lung disease in anti-MDA5-positive DM and calcinosis in juvenile DM,^30,31^ suggesting their broad pathogenic role. Fourth, we observed selective depletion of Langerhans cells in CLE, consistent with reports of reduced Langerhans cell density in non-lesional lupus skin and murine lupus models.^32^ A recent study suggested type I interferon signature has a role in Langerhans cell dysfunction, which has been proposed to lead to photosensitivity observed in lupus patients.^32^ Therefore, their loss may contribute to disease progression, as Langerhans cells also play crucial roles in maintaining skin immune homeostasis through IL-10 production and regulatory T cell activation.^33^

Patients with adult-onset dermatomyositis (DM) have a significantly elevated risk of malignancy.^4,34^ A major gap in the management of dermatomyositis is enhanced risk stratification of patients as cancer-associated versus non-cancer-associated DM at the time of diagnosis. Current guidelines for cancer screening in DM patients suggest utilizing a combination of autoantibody status and clinical features to determine risk for malignancy.^35^ Given the low threshold for a DM patient to be considered high-risk, many patients undergo extensive screening that is repeated annually for three years^35^ leading to increased health care costs, radiation exposure to patients, and substantial patient distress and/or anxiety. Therefore, understanding the underlying immune mechanisms that distinguish cancer-associated DM from non-cancer-associated DM can yield discernible markers that can enhance early risk stratification of DM, providing the opportunity for expedited detection and management of malignancies in these patients. Among DM patients, those with anti-TIF1γ have the highest risk of cancer followed by anti-NXP2.^36^ Our cohort was predominantly composed of anti-TIF1γ-positive patients and those without detectable autoantibodies, potentially influencing our observed cancer/non-cancer associations. Despite this limitation, we identified clear molecular distinctions between cancer and non-cancer DM through integrated spatial and single-cell analysis. Non-cancer DM exhibited dense immune aggregates with elevated CXCL10+ myeloid cells, neutrophils, monocytes and CXCL10+ macrophages. Moreover, these lesions were characterized by abnormal vascular endothelial and perivascular stromal remodeling, leading to loss of vascular-associated fibroblast identity (marked by ABCA8/9/10 downregulation) and upregulation of hypoxia-responsive genes in both endothelial cells and fibroblasts. In contrast, cancer-associated DM displayed more dispersed immune infiltrates with preserved vascular architecture and active angiogenesis. Notably, these cancer-associated niches showed distinctive enrichment in B cells, plasma cells and CD8+ T cells, particularly in the C_cl niche subtype, which contained substantial populations of these adaptive immune cells. This enhanced presence of adaptive immune cells, especially cytotoxic CD8+ T cells alongside B cells, may reflect active anti-tumor immune responses, similar to tertiary lymphoid structures observed in cancer.^37^ This observation becomes particularly intriguing in light of the potential hypothesis that someDM cases may be fundamentally cancer-associated but do not manifest given the patient’s ability to mount a successful early anti-tumor response.^38^ While our findings of organized B cell and CD8+ T cell responses in cancer-associated DM might support this adaptive anti-tumor immunity framework, the distinct immune architecture in non-cancer DM suggests an alternative pathway. The predominance of neutrophils and CXCL10+ monocytes in non-cancer DM, coupled with hypoxic tissue remodeling, points to a heightened innate immune response that might be more effective at early tumor elimination, albeit at the cost of tissue damage. This finding is surprising given that vasculopathic skin changes (e.g. cutaneous necrosis, cutaneous vasculitis are associated with elevated risk of cancer in DM patients)^39^. One potential explanation is that some of the TIF1γ-positive, non-cancer associated DM patients in our cohort may have mounted a vigorous immune response that led to successful elimination of cancer cells and prevented the emergence of clinically-detectable cancer. This is supported by a recent study^40^ that showed increased diversity of immune response in TIF1γ-positive DM patients is associated with decreased risk of cancer emergence. These fundamentally different immune niches and molecular programs suggest distinct pathogenic mechanisms, challenging the notion that non-cancer DM simply represents resolved cancer-associated disease.

Moreover, in our analysis we found that immune regulation is also quite distinct between cancer and non-cancer DM. In addition to a significantly higher number of CXCL10+ myeloid cells, in non-cancer DM immune niches specific cellular pairs appeared: PD-L1-expressing mregDCs and activated Tregs expressing NFKB2 and TNF receptors. In contrast, cancer-associated DM immune niches showed lower Treg abundance and lacked both PD-L1 expression and mregDC-Treg colocalization. This specific colocalization of PD-L1 expressing mregDC and activated Treg represents a surprising observation: despite the presence of this typical immunosuppressive interaction^41–43^, autoimmune inflammation still occurs. Interestingly, it has been repeatedly shown in cancer research that this mregDC-Treg pair is the cause of failure in the application of immunotherapy; this pair creates an immunosuppressive microenvironment that protects tumor cells from therapy.^44,45^ A similar pattern was observed in CLE immune niches. However, even though both mregDCs and activated Tregs were present, they also harbored substantial pro-inflammatory populations, suggesting these regulatory mechanisms may be insufficient to control autoimmunity.

Our analysis of spatial single cell data reveals new insights into the multiplicity and complexity of immune cell organization in different disease subtypes, showing the importance of further exploring spatial immune interactions in the study of skin autoimmune disease pathogenesis.

## Acknowledgements

We thank members of the Wei lab, Rashighi lab, and Korsunsky lab for helpful discussions. We thank members of the 10X Genomics team, including Rachel Gate, Steve Kujawa, Navid Farahani, Anne Monahan, Adam Semple, for their time and generous support of this project. This work is supported by a Women’s Hospital Department of Medicine - Broad Institution collaborative research Award and a Dermatology Foundation (Patient-Directed Investigation Grant). K.W. is supported by a NIH-NIAMS K08AR077037, a Doris Duke Foundation Clinical Scientist Development Award, a Burroughs Wellcome Fund Career Awards for Medical Scientists.

## Author contributions

Conceptualization: K.S.A., N.S., M.R.,I.K., R.G., R.A.V., and K.W. Single cell and spatial single cell data generation: C.G., J.L., S.P. Data analysis: K.S.A. Skin biopsy acquisition: N.S., R.L.C.,K.A., E.T., A.L., K.H. Validation: S.K. Figure generation: K.S.A., A.N.K. Writing – original draft: K.S.A.

Draft reviewing and editing: all. Supervision: K.W. Funding acquisition: K.W. All authors have read and agreed to the published version of the manuscript.

## Availability of data and materials

Further information and requests for code and data should be directed to and will be fulfilled by the Lead Contact, Kevin Wei

---

**Extended Data Fig. 1.**
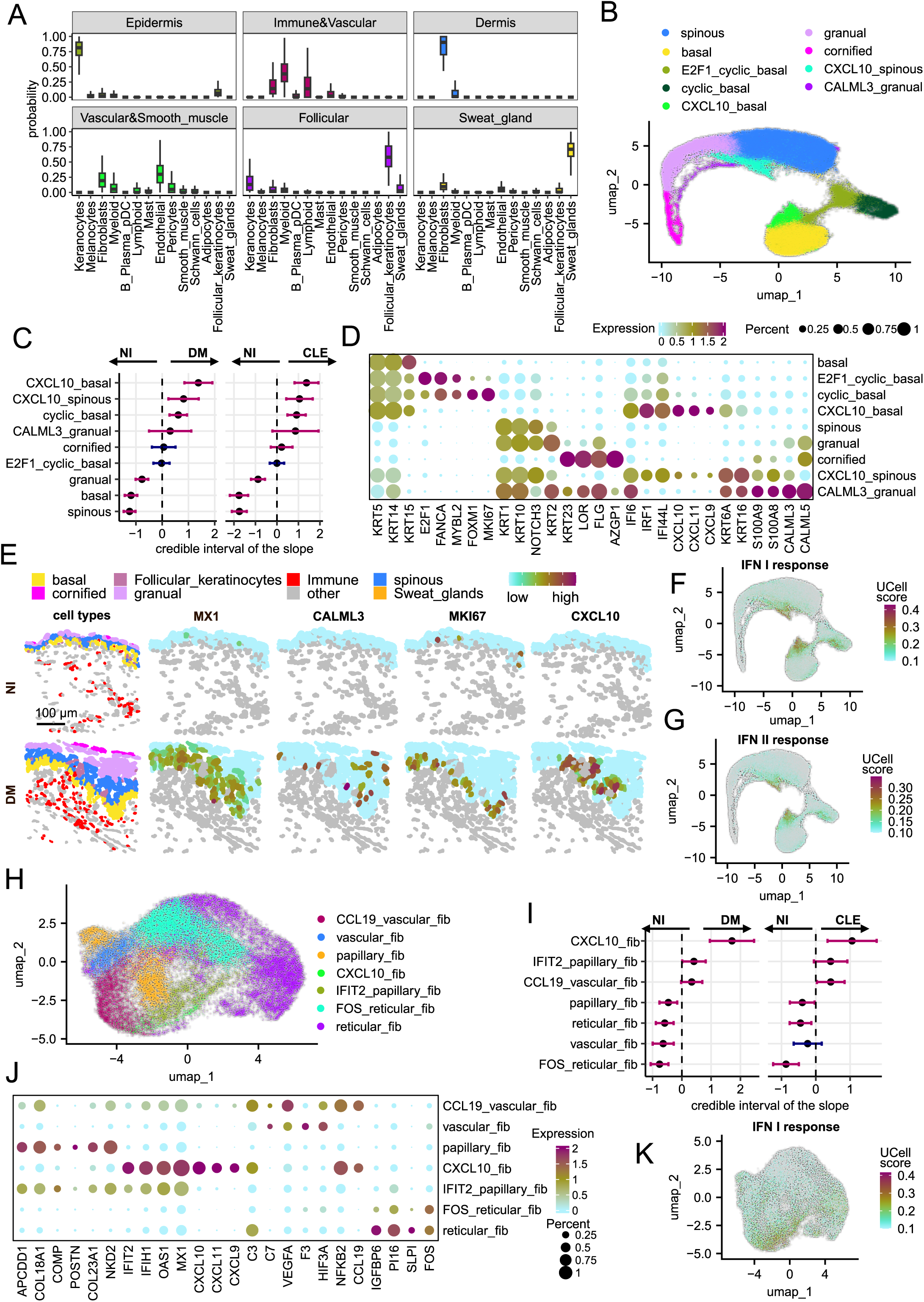
Cell state-specific analysis of interferon responses in keratinocytes and fibroblasts. **A.** Probability of spatial colocalization between each major compartment and all identified major cell states. Y-axis shows probability scores (0-1.00), with error bars indicating confidence intervals **B.** UMAP visualization of keratinocyte cell states identified by analyzing single cell data. Each point represents a single cell, colored by cell state identity as shown in the legend. **C.** Differential composition analysis of keratinocyte cell states between DM, CLE lesional and NI regions. Points show the mean posterior distribution of composition differences, with error bars representing credible intervals (2.5% and 97.5% quantiles). Red indicates significant changes (FDR < 0.1) and blue shows non-significant differences in cell state proportions. Arrows indicate which condition shows enrichment for each cell state in the respective comparison. **D.** Characterization of keratinocyte cell states by gene expression of marker genes. Dot color represents average expression level, dot size indicates percentage of expressing cells. **E.** Spatial localization of chosen cell states and their gene expression in representative NI and DM skin sections. The first column shows the distribution of cell states (color-coded according to legend). Subsequent columns display expression patterns of MX1, CALM3, MKI67, and CXCL10, with color intensity indicating expression level. Gray indicates cells with zero gene expression. **F-G**. Distribution of UCell scores for type I (F) and type II (G) interferon response gene signatures projected onto UMAP embedding of keratinocyte cell states. **H.** UMAP visualization of fibroblast cell states identified by analyzing single cell data. Each point represents a single cell, colored by cell state identity as shown in the legend. **I.** Differential composition analysis of fibroblast cell states between DM, CLE lesional and NI regions. Points show the mean posterior distribution of composition differences, with error bars representing credible intervals (2.5% and 97.5% quantiles). Red indicates significant changes (FDR < 0.1) and blue shows non-significant differences in cell state proportions. Arrows indicate which condition shows enrichment for each cell state in the respective comparison. **J.** Characterization of fibroblast cell states by gene expression of marker genes. Dot color represents average expression level, dot size indicates percentage of expressing cells. **K.** Distribution of UCell scores for type I interferon response gene signature projected onto UMAP embedding of fibroblast cell states.

**Extended Data Fig. 2.**
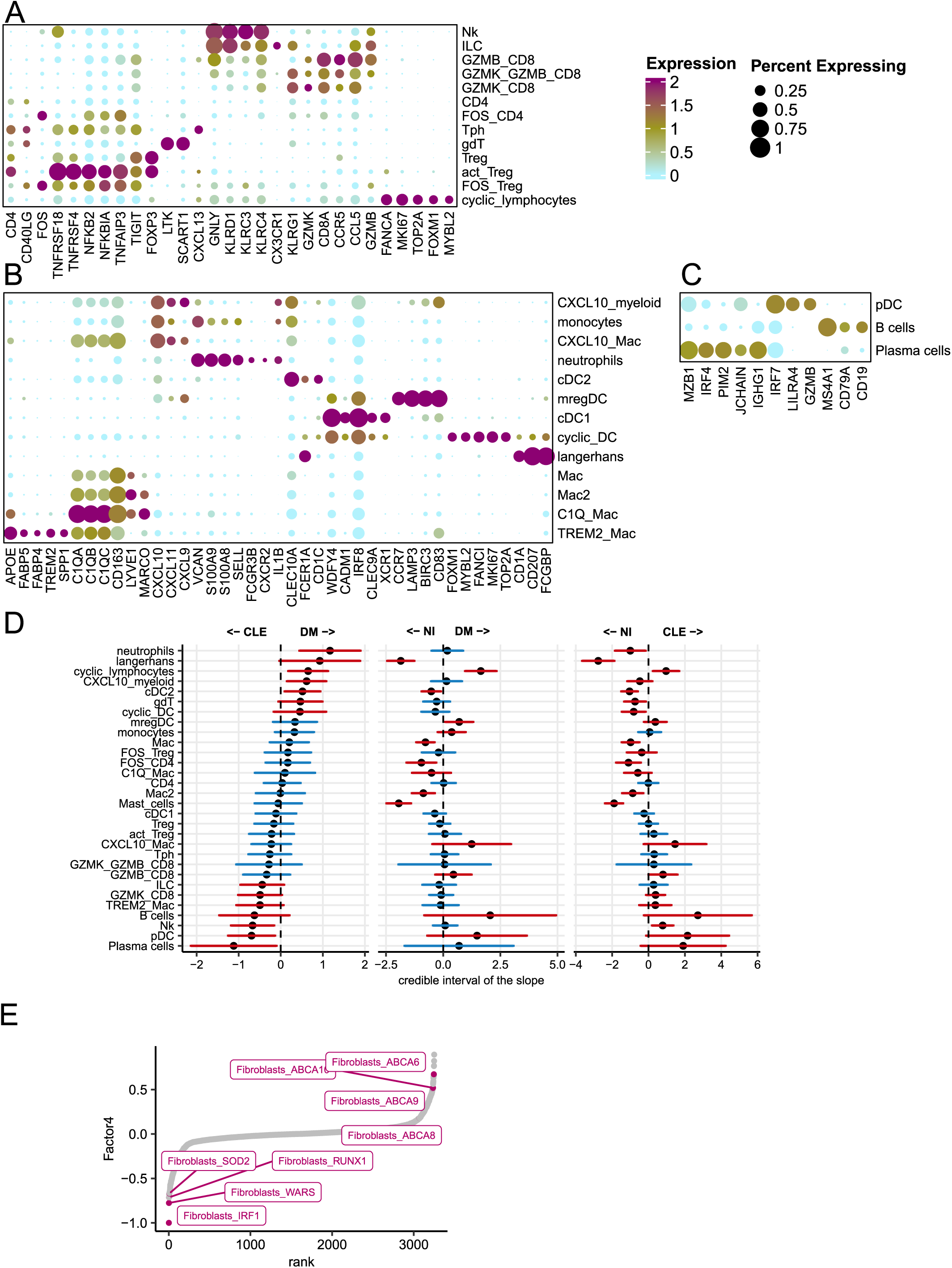
Immune cell characterization and distribution across OM and CLE. **A.** Characterization of lymphoid cell states by gene expression of marker genes. Dot color represents average expression level, dot size indicates percentage of expressing cells. **B.** Characterization of myeloid states by gene expression of marker genes. Dot color represents average expression level, dot size indicates percentage of expressing cells. **C.** Characterization of B/plasma/pDC cell states by gene expression of marker genes. Dot color represents average expression level, dot size indicates percentage of expressing cells. **D.** Differential composition analysis of immune cell states between DM, CLE and non-inflamed regions. Points show the mean posterior distribution of composition differences, with error bars representing credible intervals (2.5% and 97.5% quantiles). Red indicates significant changes (FDR < 0.1) and blue shows non-significant differences in cell state proportions. Arrows indicate which condition shows enrichment for each cell state in the respective comparison. **E.** MOFA Factor 4 loadings ranked by value. The plot displays gene weights contributing to Factor 4, with genes ranked by their contribution (x-axis) against their loading values (y-axis).

**Extended Data Fig. 3.**
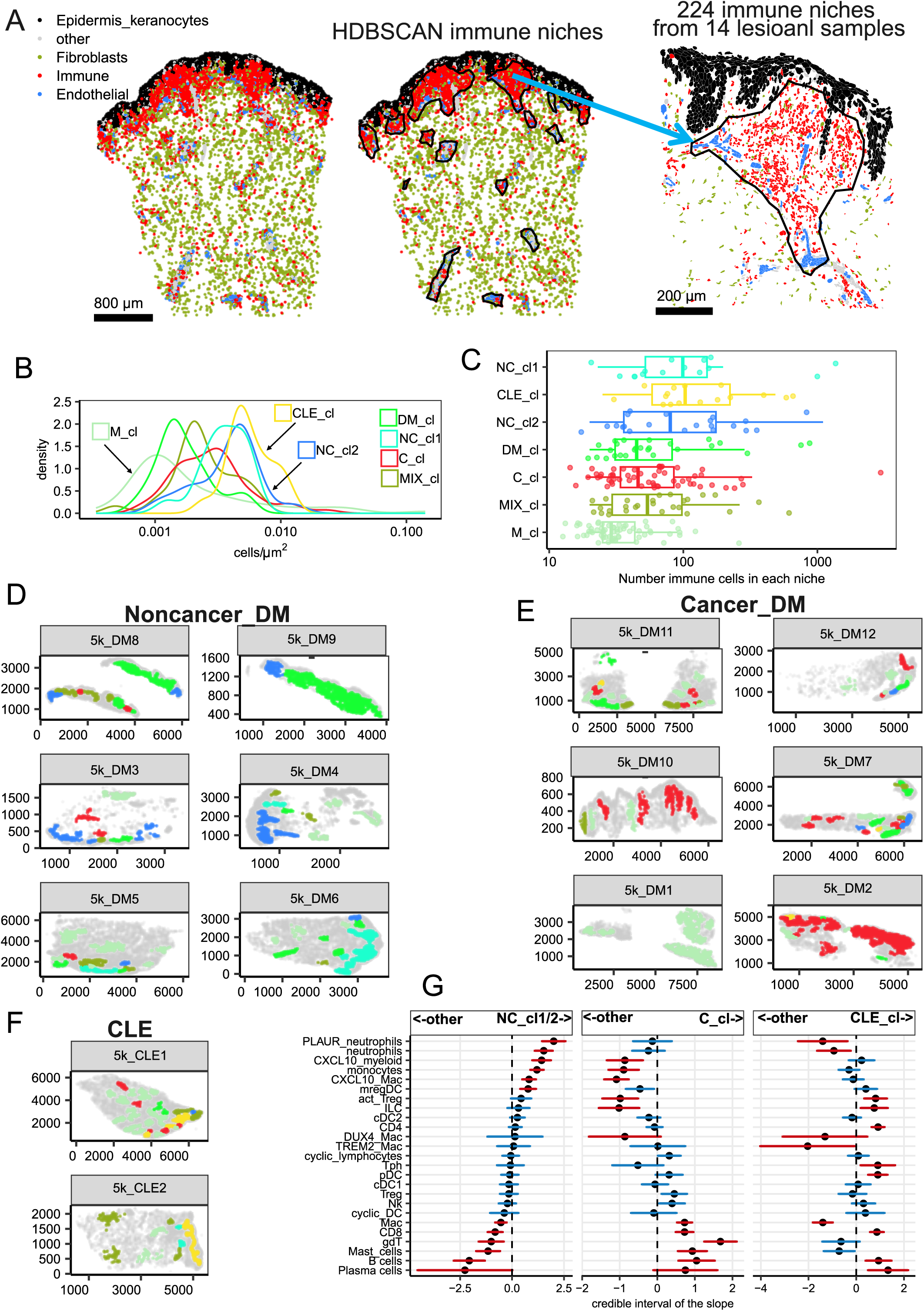
Spatial organization of immune niches distinguishes cancer and non-cancer associated OM. **A.** Pipeline for immune niche identification using spatial single cell data. Left panel: cellular composition of a representative skin section showing epidermal keratinocytes (black), fibroblasts (green), immune cells (red), endothelial cells (blue), and other cell types (gray). Middle panel: identified immune niches using HDBSCAN (black outlines). Right panel: example of the representative immune niche. **B.** Distribution of immune cell density across different niche types. The plot shows density curves of immune cell concentration (cells/μm²) for each niche type (M_cl, DM_cl, NC_cl1, NC_cl2, C_cl, CLE_cl, MIX_cl). Each curve represents the distribution of cell densities observed across all samples for a specific niche type, with colors matching the niche classification scheme used in Fig. 3A. **C.** Distribution of immune cell numbers across different niche types. Box plots show the number of immune cells per niche (x-axis, log scale) for each niche type (y-axis). Individual points represent individual niches, colored according to the classification scheme used in Fig. 3A. **D-F.** Spatial localization of immune niches across all samples analyzed through spatial single cell data. (B) Maps of immune niches in Non-cancer DM samples and (C) Cancer-associated DM samples, (D) CLE samples. Each subplot represents an individual patient sample labeled with its identifier. Niches are colored according to their classification as defined in Fig. 3A (C_cl, CLE_cl, DM_cl, M_cl, MIX_cl, NC_cl1, NC_cl2), with matching colors between panels. Axes show spatial coordinates. **G.** Differential composition analysis of immune cell states between specific niches and all other niches. Points show the mean posterior distribution of composition differences, with error bars representing credible intervals (2.5% and 97.5% quantiles). Red indicates significant changes (FDR < 0.1) and blue shows non-significant differences in cell state proportions. Comparisons are shown between: NC_cl1/2 vs other niches (left), C_cl vs other niches (middle), and CLE_cl vs other niches (right).

**Extended Data Fig. 4:**
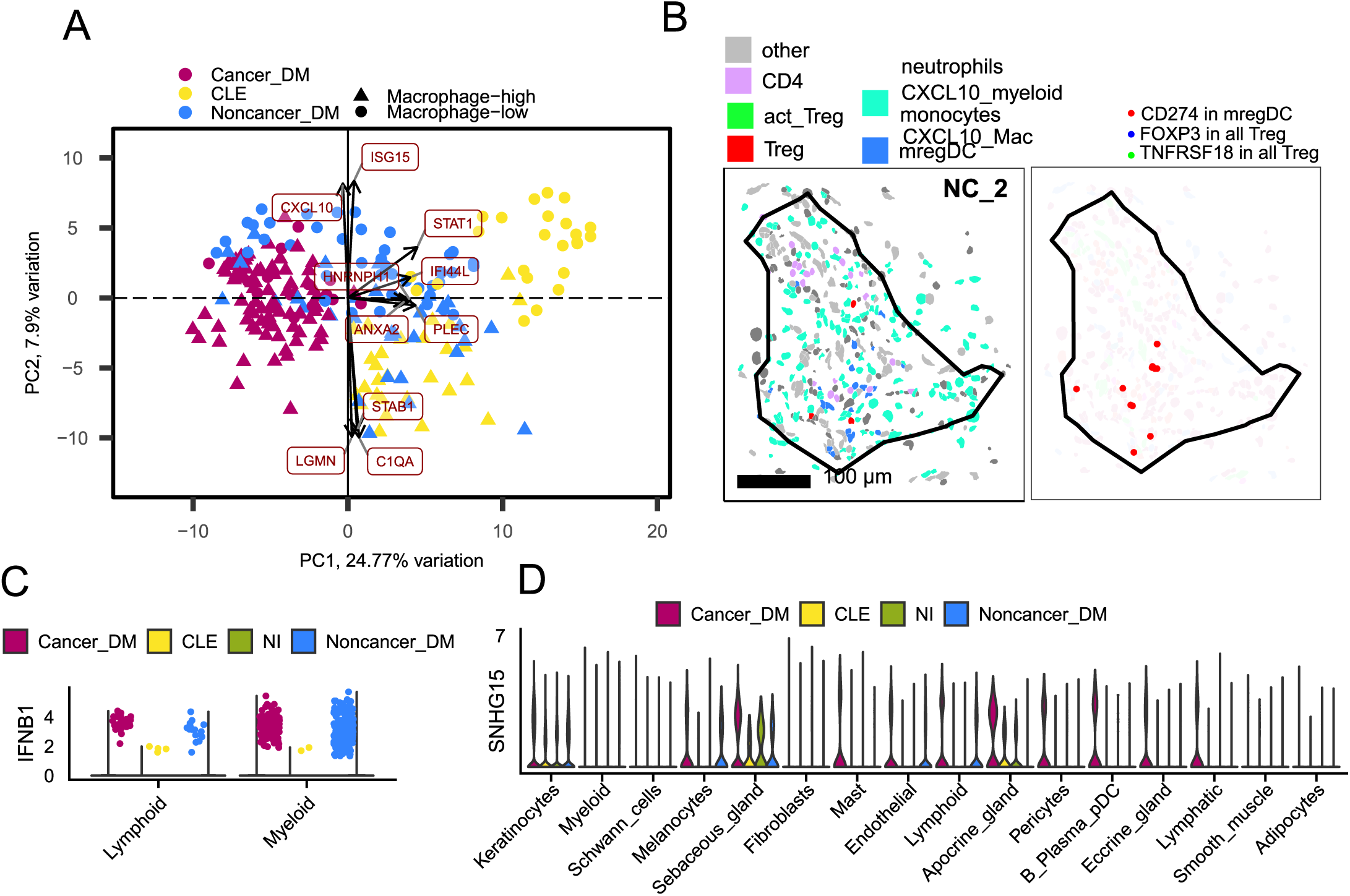
Immune niche specific transcriptomic signatures. **A.** Principal Component Analysis of pseudobulk RNA-seq data from immune niches. Each point represents an immune niche colored by disease condition: Cancer_DM (red), CLE (yellow), and Noncancer_DM (blue). Point shapes indicate niche classification based on macrophage content (triangles: Macrophage-high, circles: Macrophage-low). Key genes driving the separation are labeled. **B.** Spatial analysis of mregDC and Treg cell interactions in NC_cl2 immune niche. The left panel shows the distribution of immune cell state, colored according to the legend. The right panel displays the spatial distribution of specific transcripts: CD274 in mregDC (red), FOXP3 in all Tregs (blue), and TNFRSF18 in all Tregs (green). The niche shown is the same as in Fig. 3F, with black outlines delineating niche boundaries. **C.** Expression levels of *IFNB1* from spatial transcriptomics data across lymphoid and myeloid cell states, stratified by disease conditions: Cancer_DM (red), CLE (yellow), NI (green), and Noncancer_DM (blue). **D.** Violin plot showing the expression levels of *SNHG15* across various cell types, stratified by disease conditions through analysis of spatial single cell data: Cancer_DM (red), CLE (yellow), NI (green), and Noncancer_DM (blue).

**Extended Data Fig. 5:**
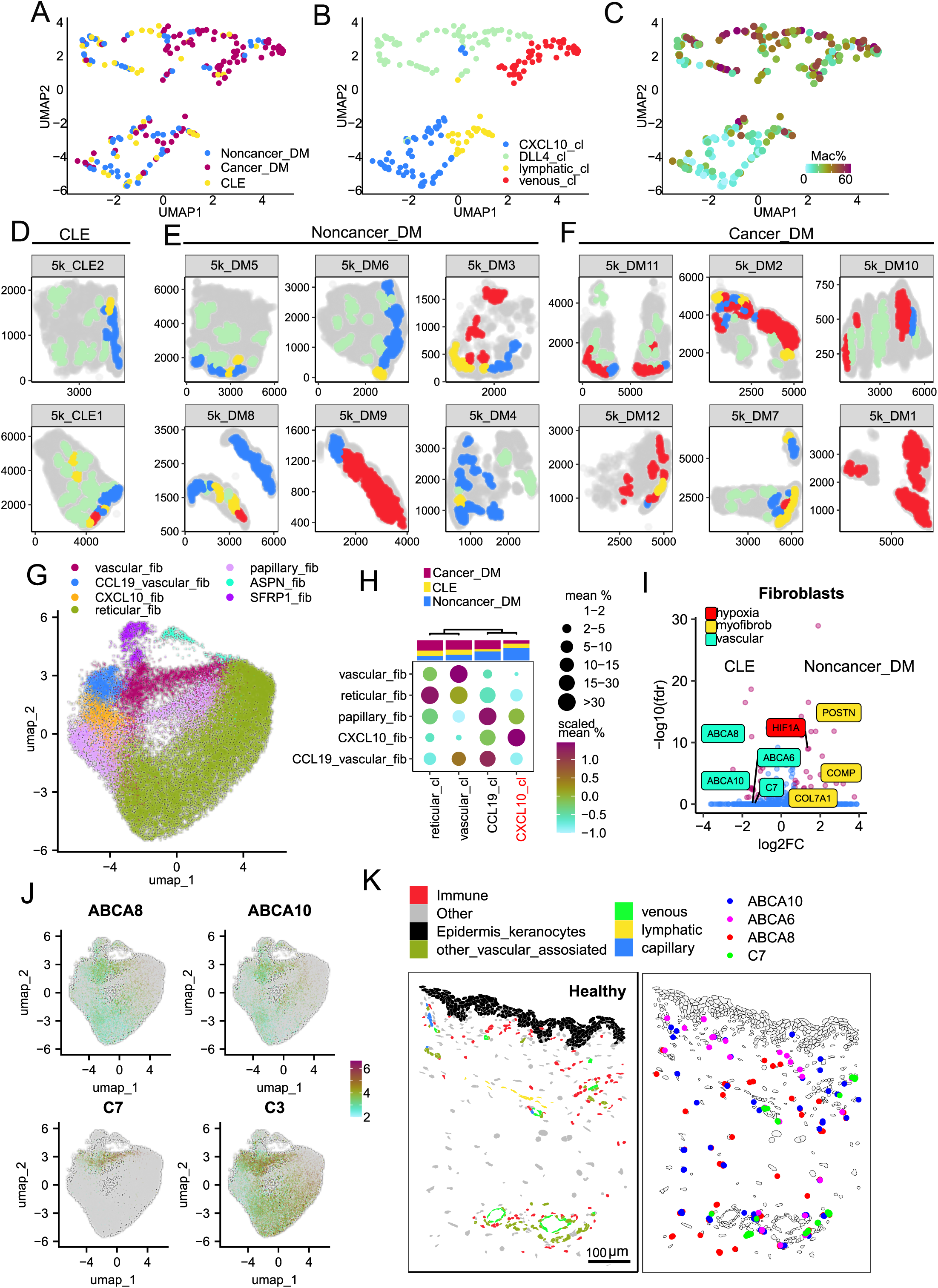
Spatial characterization of stromal and endothelial cell niches. **A-B.** UMAP visualization of endothelial cell composition within vascular niches from 14 lesional samples. (A) Left panel shows vascular niches colored by disease condition: Cancer_DM (red), CLE (yellow), and Noncancer_DM (blue). (B) Right panel displays the same aggregates colored by vascular niche type (DLL4_cl, CXCL10_cl, lymphatic_cl, venous_cl). **C.** UMAP visualization of endothelial cell composition within vascular niches, with color indicating the macrophage percentage in the originating immune aggregates. **D-F.** Spatial distribution of vascular niches across all analyzed samples. (D) Maps of immune niches in Non-cancer DM samples and (F) Cancer-associated DM samples, (E) CLE samples. Each subplot represents an individual patient sample labeled with its identifier. Niches are colored according to their classification as defined in Extended Data Fig. 5A (DLL4_cl, CXCL10_cl, lymphatic_cl, venous_cl), with matching colors between panels. Axes show spatial coordinates. **G.** UMAP visualization of fibroblast cell states identified by analysing spatial transcriptomics data. Each point represents a single cell, colored by cell state identity as shown in the legend. **H.** Fibroblasts cell composition within and adjacent to immune niches, grouped into distinct fibroblasts’ niches. The dot plot displays fibroblast states (y-axis) within each identified fibroblast niche type, where dot size represents the mean cell type percentage (ranging from 0 to >30%). Color scale indicates the relative abundance of each cell type (red: high, cyan: low). The top bar shows the distribution of samples from different conditions within each niche type: Cancer_DM (red), CLE (yellow), and Noncancer_DM (blue). **I.** Differential gene expression in fibroblasts between CLE and Noncancer_DM immune niches. Volcano plot shows gene expression differences, with significantly differentially expressed genes (|log2FC| > 0.7, FDR < 0.1; red dots) labeled and colored by pathway: fibrosis (yellow), hypoxia (red), inflammation (green), and vascular-associated signature (turquoise). Blue dots represent non-significant changes. **J.** Feature plot of vascular-associated fibroblast markers in fibroblast cell states identified through spatial single cell data analysis. **K.** Spatial distribution of vascular-associated gene signature in healthy samples. Representative tissue sections showing expression of ABCA10 (blue), ABCA8 (red), ABCA6 (pink), and C7 (green) transcripts in fibroblasts.

## Extended Data Tables

**Extended Data Table 1** - Metadata description of single-cell and spatial single-cell transcriptomics datasets.

## METHODS

### Patient recruitment and tissue sample collection

Skin tissue for analysis was derived from three independent sources–Brigham and Women’s Hospital (BWH), University of Massachusetts Memorial Medical Center (UMASS), and BWH Pathology Archies, enhancing the external validity. Fresh skin tissue acquisition was obtained through patient recruitment and sampling performed in autoimmune skin disease clinics and general dermatology clinics at BWH and UMASS. Patients were recruited with active DM and CLE (all subtypes) to provide lesional and non-lesional skin samples, and healthy patients were included to provide control tissue. Next, consent was obtained. Regions of interest on the skin were marked in preparation for tissue acquisition. Then, 1% lidocaine with epinephrine was injected into the skin to provide local anesthesia. A 4-millimeter skin punch was utilized to core out skin tissue of interest and subsequently placed in formalin. Hemostasis was achieved through suture placement. The formalin-fixed samples were then embedded in paraffin for subsequent analysis. DM and CLE formalin-fixed paraffin-embedded (FFPE) skin tissue in the BWH Pathology Archives collected within the past 12 years were also included for analysis. Samples were localized using the search terms “dermatomyositis”, “lupus”, and “interface dermatitis”. Each case was analyzed by two certified dermatologists (N.S. and R.A.V.) to ensure patient histology was concordant with a clinical diagnosis of DM or CLE before inclusion.

### FFPE single cell RNA sequencing

For each sample, four formalin-fixed paraffin-embedded (FFPE) curls (25 μm each) were dissociated following the FLEX protocol (CG000632 RevA, 10X Genomics) with the Miltenyi Biotech FFPE Tissue Dissociation Kit. Cells from each sample were hybridized with a unique Probe Barcode, as per the instructions in the "Chromium Fixed RNA Profiling Reagent Kits for Multiplexed Samples" user guide (CG000527, 10X Genomics). Post-hybridization, cells from 16 samples were washed, counted, and pooled in equal proportions. Approximately 200,000 cells from the pooled suspension were loaded onto a Chromium Q chip (PN-1000422, 10X Genomics). Sequencing libraries were prepared and sequenced on an Illumina NovaSeq platform using paired-end dual-indexing (28 cycles for Read 1, 10 cycles for i7, 10 cycles for i5, and 90 cycles for Read 2).

### Xenium slide preparation

FFPE blocks were sliced onto Xenium slides according to the “Xenium In Situ for FFPE-Tissue Preparation Guide” (CG000578 Rev C, 10X Genomics) protocol. The Xenium slides were then prepared following the “Xenium In Situ for FFPE-Deparaffinization and Decrosslinking” protocol (CG000580 Rev D, 10X Genomics). In brief, the slides were baked at 60°C for 30min and then sequentially immersed in xylene, ethanol, and nuclease-free water to deparaffinize and rehydrate the tissue. Immediately after, the Xenium slides were incubated in the decrosslinking and permeabilization solution at 80°C for 30 minutes, followed by a wash with PBS-T.

Next, the slides were prepared according to the "Xenium Prime In Situ Gene Expression" user guide (CG000760 Rev A, 10X Genomics) for the remaining steps. Probes from the Xenium Prime 5K Human Pan Tissue & Pathways Panel (PN-1000671, 10X Genomics) and the custom add-on panel were hybridized to the samples at 50°C for 18 hours. Post-hybridization, the slides underwent washing, incubation with a ligation reaction mix, another wash step, and DNA amplification. Finally, cell segmentation staining was conducted, and the slides were treated with an autofluorescence quencher and DAPI before being loaded into the Xenium instrument.

### Xenium Analyzer setup and data acquisition

Processed Xenium slides were loaded in the Xenium Analyzer and imaged, following the guidelines in the “Xenium Analyzer User Guide (CG000584 Rev F, 10X Genomics)”. After scanning, the Xenium slides were removed from the Xenium Analyzer, and processed with post-run H&E staining according to the “Xenium In Situ Gene Expression - Post-Xenium Analyzer H&E Staining” protocol (CG000613 Rev A, 10X Genomics).

### Single-cell and spatial single cell data preprocessing

The sequencing data was demultiplexed using bcl2fastq (Illumina). The resulting FASTQ files were processed with Cell Ranger using the multi pipeline and the GRCh38-2020-A reference genome. Data analysis was performed using the Seurat^46^ v5 R package. Quality control and preprocessing steps included doublet removal using scDblFinder^47^ R package and ambient RNA correction using SoupX^46,48^ R package for each sample individually. For downstream analysis, we utilized Seurat v5’s functions by splitting the data by sample, which created separate layers for each sample in a single Seurat object. This approach enabled sample-specific normalization, identification of 2000 variable features per sample, and scaling. We then performed PCA with 20 principal components and applied Harmony batch correction (lambda=1) using sample as covariate using harmony^49^ R package. Using these batch-corrected embeddings, we constructed a shared nearest neighbor (SNN) graph based on the first 20 Harmony dimensions and identified cell clusters with the Leiden algorithm (resolution 1.2). Cell type annotation was performed using established marker genes, resulting in the identification of 16 major cell states.

For spatial single-cell data, we followed a similar analytical pipeline but omitted the scDblFinder and SoupX processing steps. The only notable difference was in the normalization step, where we applied a scale factor of 5000 during normalization to account for the specific sequencing depth and transcript detection characteristics of spatial transcriptomics data.

### Major and granular cell states identification

Major cell states were first manually classified into 16 major states based on expression of known literature markers^50–52^: Keratinocytes (KRT1, KRT5, KRT17, FLG), fibroblasts (COL6A1, FBN1, PDGFRB, PI16), eccrine glands (AQP5, AQP7), apocrine glands (KRT77, GABRP), sebaceous glands (MGST1, PLIN2, FASN), melanocytes (MLANA, DST), myeloid cells (CD163, C1QA, CD74), lymphoid cells (CD3E, CD8A, CD4), B cells/plasma cells/pDC (CD19, SELL, IRF8, MZB1), Schwann cells (MPZ, SCN7A), adipocytes (PLIN1, PLIN4, ADIPOQ), mast cells (GATA2, KIT, MS4A2), vascular endothelial cells (PECAM1, VWF), smooth muscle cells (MYH11, TAGLN), pericytes (RGS5, TAGLN), and lymphatic endothelial cells (FLT4, CCL21).

For fine-grained cell state annotation, we employed the same computational approach used for major cell states but applied it to each individual cell population separately. We maintained the same parameters for normalization, variable feature identification (2000 genes), dimensionality reduction (20 PCs), and Harmony batch correction. For clustering, we adjusted the Leiden algorithm resolution parameters (1.0-1.5) to better capture the finer distinctions within each major cell type. During this process, we identified cells in the spatial data expressing markers from multiple cell populations, which we manually classified as mis-segmented and removed to define cleaner cell states.

We identified multiple granular cell states within each major cell state based on their marker gene expression profiles. For clusters without established nomenclature, we assigned names according to one of their specifically expressed markers. Cluster-specific marker genes were determined using the presto R package with an area under the curve (AUC) threshold of 0.6 ^53^.

Keratinocytes were classified into several states (**Extended Data Fig. 1D**): basal keratinocytes expressed KRT14, KRT15, and KRT5. E2F1_cyclic_basal cells showed additional expression of E2F1 with proliferation markers (MKI67, TOP2A, FOXM1), while cyclic_basal cells expressed similar proliferation markers without E2F1. CXCL10_basal and CXCL10_spinous states expressed interferon-response genes (CXCL10, CXCL9, CXCL11, IFI6, IRF1, IFI44L) at different differentiation stages. More differentiated states included spinous cells (KRT10, KRT1, NOTCH3), granular cells (KRT2, KRT23), cornified cells (LOR, FLG, AZGP1), and CALM3_granular cells characterized by calcium-binding proteins (CALM3, CALM5) alongside KRT2 and KRT23.

Fibroblast cell states(**Extended Data Fig. 1J**) included papillary fibroblasts (APCDD1, NKD2, COMP, POSTN), vascular fibroblasts (C7, C3, VEGFA, F3, HIF3A), and CCL19_vascular_fib with additional CCL19 expression. CXCL10_fib expressed interferon-response genes (CXCL10, CXCL9, CXCL11, MX1, OAS1, IFIT2), while IFIT2_papillary_fib showed interferon-response genes (MX1, OAS1, IFIT2) without CXCL10 along with papillary markers. Reticular fibroblasts expressed IGFBP6 and PI16, with a FOS_reticular subset expressing the FOS gene. Spatial transcriptomics exclusively revealed ASPN_fib (COCH, ASPN) and SFRP1_fib (SFRP1) cell states adjacent to smooth muscle cells.

Myeloid cells comprised various macrophage subsets(**Extended Data Fig. 2B**): those expressing C1QA, CD163, and MRC1; Mac2 (additional LYVE1, MARCO); C1Q_Mac (high C1QA, C1QB, C1QC); CXCL10_Mac (chemokines CXCL9/10/11); and TREM2_Mac (TREM2, APOE, SPP1, FABP4, FABP5). Dendritic cell populations included cDC2 (CD1C, CLEC10A, FCER1A), cDC1 (CLEC9A, XCR1, BATF3), mregDC (LAMP3, BIRC3), and Langerhans cells (CD1A, CD207, FCGBP). Other myeloid populations included neutrophils (FCGR3B, CXCR2, S100A9, S100A8, IL1B), with a PLAUR+ subset in spatial data; monocytes (CX3CR1, VCAN, S100A9, S100A8, CCR2); CXCL10_myeloid cells; and cyclic_DC (proliferation markers with DC markers).

Endothelial cells included tip cells (PGF, CXCR4), venous endothelial cells (CLU, VWF, CCL14), arterial endothelial cells (SEMA3G, ELN, SULF1, HEY1), and capillary endothelial subtypes (CD36, RGCC): FABP4_capillary (FABP4) and HOXD10_capillary (HOXD10). Spatial data revealed DLL4-expressing variants (DLL4_venous, DLL4_lymphatic, DLL4_artery, DLL4_FABP4_capillary) and interferon-responsive subtypes (CXCL10_venous, CXCL10_capillary). Lymphatic endothelial cells expressed FLT4, SEMA3A, and FOXC2, while cyclic_vascular cells showed proliferation markers MKI67 and TOP2A.

Lymphocyte populations (**Extended Data Fig. 2A**) included CD4+ T cells (CD4, CD40LG) with a FOS_CD4 expressing the FOS gene; T peripheral helper cells (Tph, CXCL13+); Tregs (FOXP3, TIGIT) with act_Treg (TNFRSF4, NFKB2, TNFRSF18, NFKBIA) and FOS_Treg variants; gamma-delta T cells (LTK, SCART1); and cyclic_lymphocytes (MKI67, TOP2A). CD8+ T cell subsets were distinguished by varying gene expression levels of GZMB, GZMK, and CD8A, resulting in three populations: GZMK_GZMB_CD8, GZMB_CD8, and GZMK_CD8. Natural killer cells expressed GNLY, KLRD1, KLRC3, and KLRC4, while innate lymphoid cells showed NK markers with CX3CR1 and KLRG1.

B cells (**Extended Data Fig. 2C**) expressed CD19, plasma cells showed high MZB1 expression, and plasmacytoid dendritic cells were characterized by IRF7, LILRA4, TCF4, and GZMB expression.

### Spatial analysis of cell states organization - compartment analysis

For global tissue organization and skin compartment identification, we employed the hoodscanR^13^ R package to analyze cellular neighborhoods based on spatial proximity. This approach involved a two-step process: first, we identified k=100 nearest neighboring cells for each cell and calculated neighbor association probabilities; second, we merged these probabilities by cell type and assessed neighborhood colocalization patterns using Pearson correlation of probability distributions between different cell states. Based on these probability distributions, we applied k-means clustering with 6 centers to define distinct tissue compartments.

### Cell composition analysis

To visualize differences in cellular composition between samples, we first calculated the percentage of each cell type per sample. We then used Bray-Curtis dissimilarity to measure how different samples were from each other based on their cell type proportions, and visualized these relationships using Principal Coordinate Analysis (PCoA) implemented in the labdsv R package. To assess significant differences in cell type composition across groups, we employed the sccomp^12^ R package. Sccomp implements a Bayesian framework using sum-constrained independent Beta-binomial distributions to analyze differential composition. Statistical significance was determined using two criteria: (1) a 95% credible interval of the slope, where positive and negative intervals indicated cluster enrichment or depletion respectively, and (2) an FDR threshold of 0.05. The model assessed compositional changes against a fold-change threshold of -0.1 to 0.1.

### Sample stratification using Multicellular factor analysis

To investigate transcriptional programs underlying cancer-associated DM, we employed Multicellular Factor Analysis (MOFA), implemented in the MOFAcellulaR^15^ R package. MOFA is an unsupervised method that extends PCA by identifying latent factors that capture major sources of variation in gene expression across different cell types. For data preprocessing, we generated pseudo-bulk profiles by aggregating single-cell transcriptomes within each cell type and sample combination. We filtered cell states to include only those with at least 25 cells per sample(B cells, plasma cells, and pDCs, melanocytes) and excluded cell states with inconsistent representation across samples (sweet gland and hair follicular cells). Genes for MOFA analysis were filtered through multiple steps: first, retaining genes with a minimum count of 100 in at least 25% of samples. Data were then normalized using TMM with a scaling factor of 1,000,000. To minimize the impact of noisy mRNA background contamination, we implemented a strict filtering strategy for highly variable genes. For each cell type, we excluded any highly variable genes that were identified as markers for other cell types. Cell-type-specific markers were identified through a quasi-likelihood framework (FDR < 0.01, logFC > 1) implemented in edgeR^54^ R package. These filtered and normalized matrices were then converted to MOFA format using the pb_dat2MOFA function. We fit the MOFA model with eight factors using default parameters. Factor importance was quantified through variance explained (R2) across cell types. Associations between factors and clinical variables were assessed using Kruskal-Wallis test for categorical variables (disease status, gender, site of biopsy) and Pearson correlation for continuous variables (age, immune infiltration ratio), with Benjamini-Hochberg correction for multiple testing.

### Identification and analysis of immune niches in lesional samples

To identify and characterize immune cell niches within the tissue, we implemented a multi-step computational approach. First, we performed spatial clustering using HDBSCAN (implemented in dbscan R package) to identify immune cell niches. The clustering included both immune and endothelial cells since immune cells typically aggregate around blood vessels. Clusters containing fewer than 20 immune cells or dominated by vascular cells (>90% of total cells) were excluded from further analysis. For samples with technical replicates on the same slide, we first separated the replicates using k-means clustering (k=2) and then analyzed each replicate independently.

After identifying immune niches across all 14 lesional samples, we calculated the relative abundance of each immune cell state as a percentage of total immune cells within each niche. To classify distinct immune niche subtypes, we first computed Bray-Curtis dissimilarities between all niches based on their immune cell state composition. We then performed Principal Coordinate Analysis (PCoA) and used the first four components for UMAP dimensionality reduction. Finally, hierarchical clustering was applied to define immune niche subtypes.

To identify gene expression programs within immune niches, we performed pseudobulk analysis by aggregating expression counts from all immune cells within each immune niche. Prior to differential expression analysis, we filtered out lowly expressed genes (present in <10% of aggregates with counts ≥5). We then performed differential expression analysis using DESeq2^54,55^ R package across three conditions: CLE, cancer-associated DM, and non-cancer associated DM immune niches. Genes were considered differentially expressed if they met the significance threshold of adjusted p-value < 0.1 and |log2 fold change| > 0.7.

### Finding stroma associated with immune niches

After identifying immune niches, we determined their precise boundaries and associated stromal cells. For each immune aggregate, we delineated border cells using the identify_bordering_cells function from the SPIAT^56^ R package, which implements an alpha-hull algorithm. The alpha parameter, which determines the hull shape precision, was dynamically adjusted based on niche size: alpha = 60 for niches with <200 cells, alpha = 90 for >5000 cells, and scaled linearly for intermediate sizes (alpha = (n_cells - 300)/160 + 60). The algorithm first created a convex polygon around each aggregate using the point.in.polygon function from the sp R package to identify cells within and at the border of each niche. We identified two stromal cell populations around each immune aggregate: stromal cells located within the aggregate boundary and those outside the aggregate within a 50-micrometer radius. To characterize stromal niches, we applied the same pipeline used for immune niche classification.

To identify differentially expressed genes in stromal cells associated with immune aggregates, we used MAST^57^ implemented in Seurat R package. To account for sample-specific effects, we included sample identity as a latent variable in the model.

### Cell interactions analysis

To analyze local cell-type interactions, we used the crawdad^58^ R package, examining spatial relationships at multiple distances (5, 20, 50, 100, 250, 500, 700 μm). Statistical significance of cell-type associations was determined using permutation tests (n=3) with Bonferroni correction. Z-scores were calculated to measure the likelihood of cell-type co-localization, where higher Z-scores indicate stronger spatial associations between cell populations.

### Spatial correlation gene expression analysis

To identify gene expression patterns within immune aggregates and their surrounding 50-micrometer border regions, we employed InSituCor^59^ R package. For each cell, we generated an environment expression profile by averaging gene expression across its 50 nearest neighbors. To control for potential confounding factors such as cell type abundance and transcript count, we calculated a conditional correlation matrix from residuals after regressing the environment expression against these confounders.

### Quantification of endothelial cell density in standardized tissue regions

To define regions for endothelial density analysis, we created interactive polygons that extended 600 μm from the epidermal border. Using the sf R package, we constructed precise boundary regions for each sample, allowing standardized measurement across different tissue sections. These polygons were used to identify which cells fell within the analysis regions, enabling consistent spatial quantification between samples despite variations in tissue morphology.

### Pathway and gene signature analysis

We used the decoupleR^60^ R package to estimate pathway activities across studied conditions. Pathway scores were generated using the multilevel model with the top 500 target genes per pathway. For interferon response analysis, we quantified type I and II interferon signatures from the Reactome database using UCell^61^ R package, which calculates signature enrichment in individual cells through a rank-based approach.

### Fibroblast сytokines and hypoxia treatment

Human dermal fibroblasts were seeded at a density of 2 x 10^4^ cells/well in quadruplicate in two sets of 96-well plates. Cells were cultured in complete media (5% fetal bovine serum, HEPES, MEM amino acids, L-glutamine, penicillin-streptomycin, nonessential MEM amino acids, 2-mercaptoethanol, and gentamicin). Plates were incubated overnight at 37°C in a 5% CO^2^ incubator. The next day, media was removed, and cells were treated as follows: complete condition media (control), complete media with IL1-β (1 ng/mL), complete media with IFN-β (5 ng/mL), or complete media with IFN-γ (5 ng/mL). One set of plates was transferred to a hypoxia chamber maintained at 0.5% O^2^, while the other set was maintained under normoxic conditions. After a 24-hour incubation period, RNA was extracted using TRIzol reagent according to the manufacturer’s protocol. cDNA synthesis was performed using the QuantiTect Reverse Transcription kit. Quantitative PCR (qPCR) was carried out using the Brilliant III Ultra-Fast SYBR Green qPCR master mix on an Agilent AriaMx Real-Time PCR system.

